# Ultrastructural comparison of fixation and cryopreservation methods for brain preservation

**DOI:** 10.64898/2026.09.18.751896

**Authors:** Alexander German, Cassandra Flügel-Koch, Friedrich Paulsen, Jürgen Winkler, Andrew T. McKenzie

## Abstract

Maximizing the morphological and molecular fidelity of preserved mammalian brain tissue is essential for basic neuroscience and brain banking, where tissue quality determines the reliability of downstream analyses. Aldehyde fixation and cryogenic storage are the most powerful preservation techniques available. However, how to best combine them is incompletely characterized. Here, we tested twelve different approaches to preserve the murine brain, including standard aldehyde perfusion fixation, variants of aldehyde-stabilized cryopreservation (ASC) using ethylene glycol (EG), aldehyde fixation followed by unprotected freezing, and several cryopreservation protocols without preceding aldehyde fixation, such as interleaved equilibration with vitrification solution. Preservation quality was assessed using light microscopy, transmission electron microscopy, and patch-level feature analysis with the DINOv2 vision foundation model. We found that ASC with sodium dodecyl sulfate (SDS)-mediated blood-brain barrier permeabilization preserved nuclear morphology, myelin periodicity, and neuropil texture comparable to standard aldehyde perfusion fixation, providing the first independent replication of ASC. Omitting SDS caused severe parenchymal dehydration, due to a mismatch between water and cryoprotectant transport. Aldehyde fixation followed by unprotected slow freezing confined ice damage primarily to perivascular zones, while fast freezing led to intranuclear clefts and cavities. Among non-fixation protocols, interleaved equilibration with vitrification solution preserved myelinated axon profiles and chromatin patterns, but induced perivascular edema. High concentration glycerol perfusion caused significant osmotic dehydration. Straight freezing of unfixed tissue without cryoprotectant was followed by membrane disruption, but resulted in the visualization of more structure than expected, likely as a result of structural restitution upon thawing. Taken together, our results provide a framework for matching preservation strategies to the needs of different types of brain banking and neuroscience research.

**Highlights:**

- ASC with SDS preserves brain ultrastructure as well as standard fixation.
- Omitting SDS in ASC restricts cryoprotectant entry, causing severe dehydration.
- Slow freezing of aldehyde-fixed brains limits ice damage to perivascular zones.
- Interleaved equilibration retains unfixed myelin but induces vasogenic edema.
- Unprotected freezing disrupts membranes despite partial structural restitution.

## Introduction

Brain tissue is a delicate material of high information density. Maximizing the morphological and molecular fidelity of preserved brain tissue is an important aspect of brain banking, wherein achieving high quality tissue preparation is essential to ensure the reliability of downstream analyses. A better understanding of preservation methods is also important for performing basic neuroscience research on the biomolecule-annotated connectome. Among the numerous conditions that are known to preserve brain tissue to at least some degree [1], aldehyde fixation and cold temperature are known to be the most powerful techniques [2]. Here, we explore how they may be combined to achieve morphological and molecular preservation at the same time.

Fixation with formaldehyde and/or glutaraldehyde is a standard step in processing samples for microscopy, including for connectomic studies. The chemical crosslinks introduced by these agents between primary amines are below the ultrastructural resolution limit and provide strong structural stabilization [3, 4]. Glutaraldehyde offers particularly good ultrastructural fixation, but protein crosslinking can mask epitopes and make immunocytochemical staining more difficult. Antibodies raised against glutaraldehyde-fixed antigens can improve antigen recognition in fixed tissue [5]. Aldehyde polymerization stabilizes all macromolecular tissue constituents into a gel that carries substantial information about gene expression in the native tissue [6]. Some expect that aldehyde fixation alone can allow tissue to last for up to several centuries at ambient temperature, given the slow spontaneous hydrolysis rate of the peptide bond [7]. However, aldehyde-fixed tissue stored at ambient or refrigerated temperatures undergoes progressive loss of antigenicity over months to years, limiting the window for molecular studies such as immunohistochemistry and proteomic analyses [4]. Combining aldehyde fixation with cryoprotectant loading and subzero storage offers a route to long-term preservation of both ultrastructure and molecular epitopes [8].

Preservation of all molecular properties of biological material, including labile post-translational modifications of proteins, RNA and metabolites can be achieved by lowering the temperature, as described by the Arrhenius equation [9]. Below the glass-transition temperature of water at approximately -130°C, molecular mobility becomes arrested completely, leaving ionizing radiation of cosmic and terrestrial origin as the remaining source of molecular damage [10]. The universal preserving effect of cold temperature on biomolecules is traded off against the potential for morphological displacements at the nanoscopic scale due to the reorganization of water into ice crystals. Ice formation can be modified via cooling and rewarming rates and the addition of cryoprotective agents (CPAs), which can be used to suppress ice formation entirely via vitrification [10]. The use of cryoprotectants is complicated by osmotic volume shifts that result from a difference in cell membrane permeability of cryoprotectants and water, which is exacerbated by the low permeability of the blood-brain barrier (BBB) [11].

Naturally freeze-tolerant species permit, and frequently initiate, extracellular ice formation, protecting their intracellular architecture against consequent osmotic dehydration through the accumulation of low-molecular-weight osmolytes [12].

The wood frog Rana sylvatica is the best-characterized vertebrate model of freeze tolerance, in which hepatic glycogenolysis mobilizes glucose for distribution to peripheral tissues [13]. Under these conditions, approximately two-thirds of total body water transitions into extracellular ice without compromising viability [14]. Northern Alaskan populations can withstand cooling to −16 °C, facilitated by high concentrations of cryoprotectants within brain tissue [15]. These cryoprotectants are therefore endogenous metabolites actively distributed via the amphibian circulation across the blood-brain barrier or synthesized locally.

Attempts to extend freezing tolerance to mammals were initiated by the identification of glycerol as a viable cryoprotectant for isolated cells [16]. Early on in cryobiology, it was demonstrated that hamsters could be partially frozen below 0 °C and subsequently resuscitated [17], with the observation that crystallization up to 60% of cerebral water was compatible with the return of consciousness [18].

By utilizing 15% glycerol, subsequent studies by Suda et al. achieved partial retention of bioelectric excitability in frozen feline brains [19, 20]. However, the overt mechanical damage inflicted by crystallization can be avoided in ice-free preservation via vitrification [21]. Vitrification has enabled reliable cryopreservation of structure and function in brain slices, while vascular cryoprotectant delivery and removal in the intact mammalian brain is challenging [22–24].

Aldehyde-stabilized cryopreservation (ASC) was introduced as a method to combine the benefits of both aldehyde fixation and cryopreservation [25]. By perfusing aldehyde fixatives followed by cryoprotectants and then cooling to cryogenic temperatures, ASC allows brain tissue to be stored indefinitely while potentially retaining molecular epitopes that degrade over months to years in fixative at ambient or refrigerated temperatures. Pre-fixation before cryoprotectant loading may also mitigate the osmotic damage that cryoprotectants can cause on unfixed tissue, as aldehyde crosslinking mechanically stabilizes membranes and attenuates the cell shrinkage and swelling that otherwise accompanies cryoprotectant infiltration [26]. However, effective cryoprotectant penetration into the brain parenchyma requires permeabilization of the BBB with detergents such as sodium dodecyl sulfate (SDS), which solubilizes membrane lipids and may wash out tissue components. To the best of our knowledge, a replication study of ASC has not yet been published.

Protocols using cryopreservation without aldehyde fixation are of particular interest because they are compatible with non-morphological molecular biological techniques, such as bulk RNA sequencing, requiring the dissociation of molecules, and may even allow for the read out of electrophysiological properties [19, 22]. However, they have not yet demonstrated comparable ultrastructural preservation quality in the available literature, at either the two-dimensional or volumetric level, potentially due to difficulty in loading and unloading cryoprotectant without osmotic damage [2, 10, 27].

In this study, we tested 12 different approaches to preserve the mammalian brain, including conventional aldehyde perfusion fixation (labeled conditions A and B), variants of ASC with 65% ethylene glycol (conditions C, D, E, and F), and fast and slow freezing after aldehyde fixation without cryoprotection (conditions G and H). We also tested several pure cryopreservation protocols without the use of preceding aldehyde fixatives, namely 65% glycerol as introduced in 1995 by Darwin et al. (condition I, [28]), with 15% glycerol as introduced in 1966 by Suda et al. (condition J, [19, 20]), unprotected cryopreservation (also known as “straight freezing”, condition K), and interleaved equilibration with 59% V3 vitrification solution, as introduced in 2025 by German et al. (consisting of dimethyl sulfoxide (DMSO), ethylene glycol and formamide dissolved in water, condition L, [22]). For each method, we studied the preservation quality using both light and electron microscopy.

## Methods

### Animals and ethical approval

All animal-related experimental procedures were performed according to the law of animal experimentation issued by the German Federal Government. Experimental procedures were approved by the local animal welfare authority (Regierung von Unterfranken, AZ: 55.2.2-2532-2-1109 and RUF-55.2.2-2532-2-2147-13). C57BL/6 mice were used for all initial perfusion experiments (male, age 6-9 months). Fischer 344 Laboratory rats were used for follow-up aldehyde immersion fixation experiments (male, age 12 months).

### Transcardial perfusion

Under isoflurane anesthesia, mice were sacrificed. The vascular system was cleared from blood via transcardial perfusion of 20 mL of 4°C phosphate-buffered saline (PBS, 1x) injected manually over 2 minutes with a 21G needle and syringe into the left ventricle following incision of the right atrium. PBS composition was NaCl (137 mM), KCl (2.7 mM), KH₂PO₄ (1.5 mM), Na₂HPO₄ (8.1 mM), pH 7.4.

Subsequently, the descending aorta was clamped at the level of the diaphragm with large hemostatic forceps and all caudal tissues removed. The resulting cranial cephalothoracic specimen was transferred to a 4°C PBS bath. The left ventricle was opened via resection of the cardiac apex under a stereomicroscope and a 0.8 mm diameter button cannula (Acufirm Ernst Kratz, Germany) was advanced into the aortic valve in a manner that its equator did slightly penetrate beyond the atrioventricular plane. The cannula was secured in place with small hemostatic forceps with curved serrated jaws (BH109R, Aesculap, Germany). These steps took approximately 8 minutes to complete. All subsequent perfusates were delivered via a peristaltic pump (Reglo, Ismatec, Germany) at a flowrate of 4 ml/min for non-CPA containing solutions, and 1 ml/min for CPA containing solutions to account for the increased viscosity.

### Brain preservation conditions

Twelve preservation conditions were tested (labeled A through L; **Table 1**, **Fig. 1**). All conditions began with the PBS perfusion described above. One animal was used per condition. The specific protocols were as follows:

**Figure 1.**
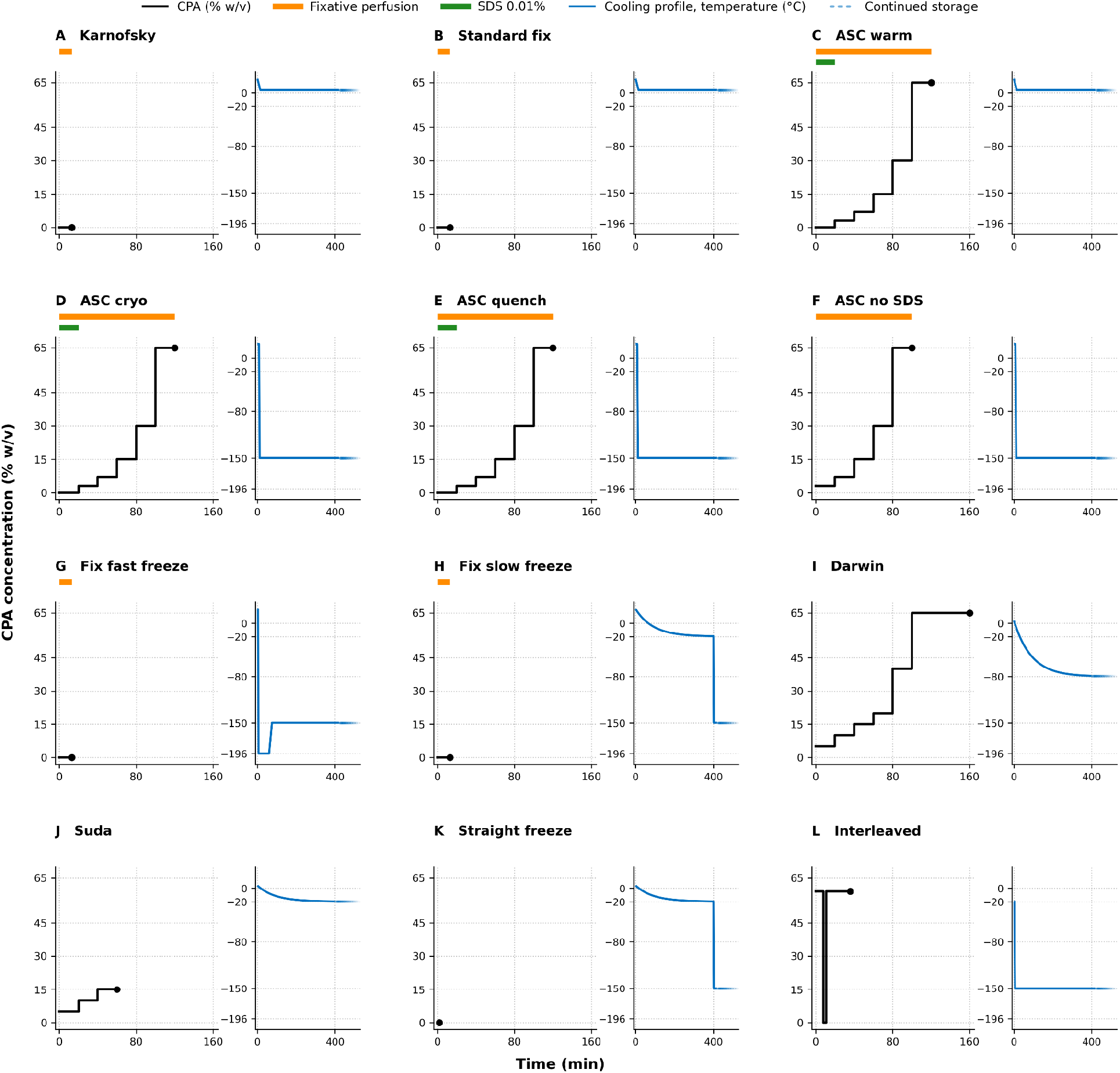
Schematic overview of the twelve brain preservation conditions. (A–L) correspond to the conditions in Table 1. For each condition, the left subpanel shows the CPA loading trajectory during perfusion (black line; ethylene glycol in C–F, glycerol in I–J, V3 vitrification solution in L), with the filled circle marking the end of the perfusion period and a flat line at 0% indicating the absence of cryoprotectant (A, B, G, H, K). Horizontal bars above each panel indicate the fixative perfusion window (orange) and the 0.01% SDS permeabilization step (green). The right subpanel shows the subsequent cooling profile to the storage temperature (blue line), drawn schematically from the protocol parameters; measured thermal profiles for slow freezing are shown in Fig. S1.

**Table 1.**
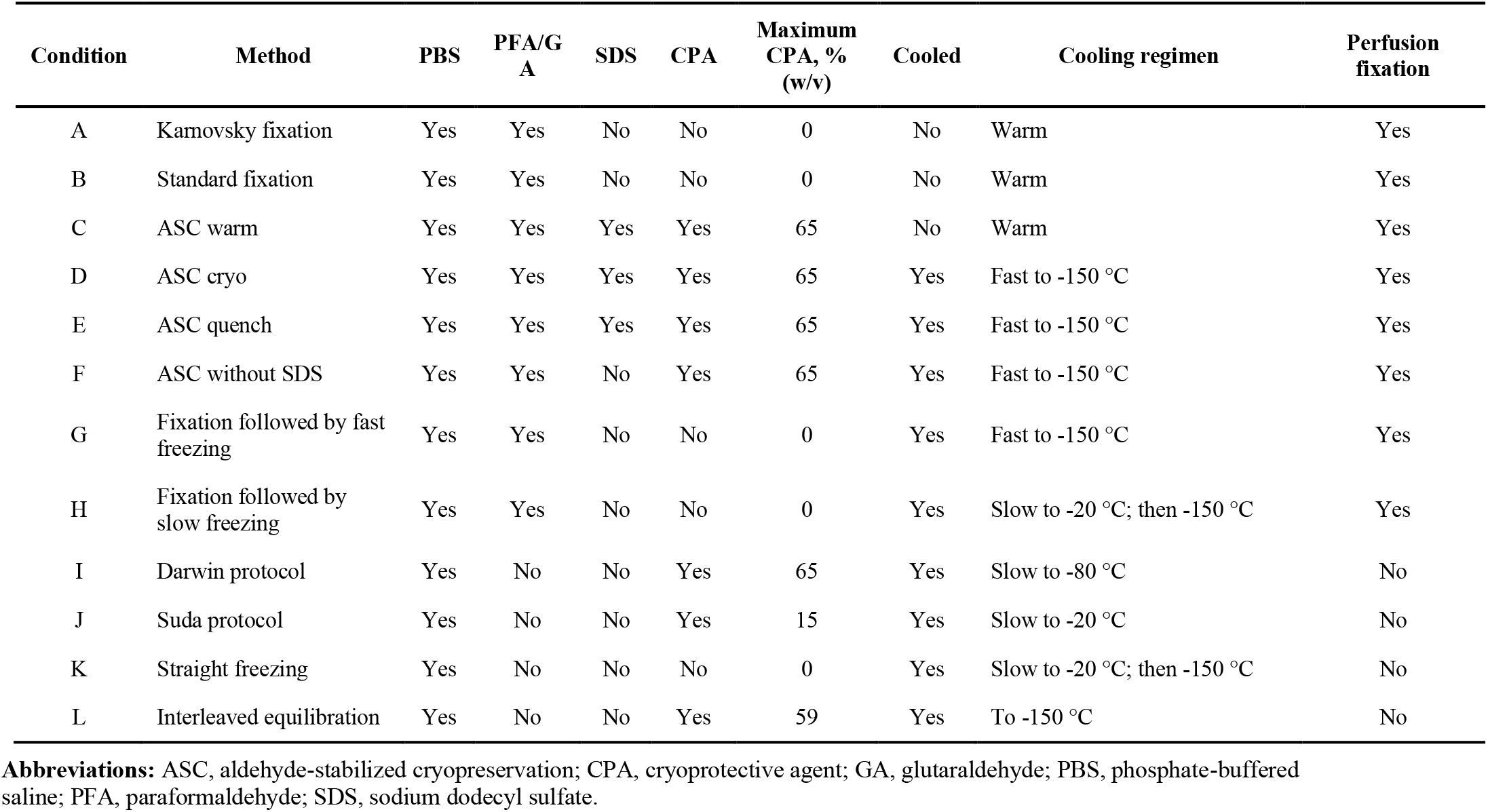
Preservation conditions and key protocol parameters.

Condition A (Karnovsky fixation): Perfusion with 50 ml of Ito/Karnovsky fixative, consisting of 2.5% paraformaldehyde (PFA) and 2.5% glutaraldehyde (GA) in 0.1 M cacodylate buffer with 1 µg/mL picric acid [29].

Condition B (Standard fixation): Perfusion with 50 ml of 1% (w/v) PFA and 2% (w/v) GA in PBS.

Condition C (ASC warm): Perfusion with condition B fixative, waiting for 4 hours at room temperature, followed by perfusion of 0.01% (w/v) sodium dodecyl sulfate (SDS) in fixative for 20 min, then a graded series of ethylene glycol (EG) at concentrations of 3%, 7%, 15%, 30%, and 65% (w/v) in fixative for 20 minutes each. Specimens were stored at 4°C immersed in CPA+fixative for 1 week prior to analogous gradual CPA washout of <1 mm^3^ tissue blocks on a shaker and storage for 1 further week in fixative at 4°C prior to Epon embedding.

Condition D (ASC cryo): Identical to condition C, followed by brain extraction, immersion in 5 ml CPA and fixative in a glass vial, cooling and storage for 1 week by placing the vial in the gas phase of a -150°C freezer, followed by rewarming at room temperature, analogous gradual CPA washout of <1 mm^3^ tissue blocks on a shaker and storage for 1 further week in fixative at 4°C prior to Epon embedding.

Condition E (ASC quench): Initial perfusion and cryoprotection is identical to condition C. Next, the vascular system was cleared by persufflation – i.e., gaseous perfusion – with 50 ml air using manual injection with a 50 ml perfusor syringe (Omnifix 4617509F, B Braun, Germany). 200 ml isopentane was solidified in liquid nitrogen, and rewarmed at room temperature. As soon as approximately 50% of the solution was melted (Tm = -160°C), the liquid phase was aspirated with the perfusor syringe and injected manually, followed by immersion in the remaining isopentane and storage for 1 week at -150°C. We call this procedure perfusion quenching [30].

Condition F (ASC without SDS): Perfusion with condition B fixative, followed by the same graded EG series as condition C (3%, 7%, 15%, 30%, 65% w/v, 20 minutes each) but without the SDS permeabilization step, then cooling and storage at -150°C as in condition D.

Condition G (Fixation followed by fast freezing): Perfusion with condition B fixative, waiting for 4 hours at room temperature, then rapid freezing via direct immersion of the whole, non-craniectomized cephalothoracic specimen in liquid nitrogen, followed by storage for 1 week at -150°C, followed by rewarming at room temperature, brain extraction and storage for 1 further week at 4°C before Epon embedding.

Condition H (Fixation followed by slow freezing): Perfusion with condition B fixative, followed by slow freezing by placing the specimen in a 50 ml Falcon tube inside a Styrofoam container at -20°C, reaching -20°C within approximately 400 minutes (thermal profile in **Fig. S1**), with subsequent transfer to -150°C storage.

Condition I (Darwin protocol, also known as high concentration glycerol): Perfusion with a graded series of glycerol in Hanks’ Balanced Salt Solution (HBSS) at 4°C, at concentrations of 5%, 10%, 15%, 20%, 40%, for 20 minutes each and 65% for 60 minutes [28]. The cephalothoracic specimens were then slowly cooled in a 50ml Falcon tube inside a Styrofoam container to -80°C. Rewarming was performed within approximately 10 minutes by stirring the Falcon tube in water at room temperature, followed by rapid brain extraction and immersion in 4% PFA fixative at 4°C for 1 week.

Condition J (Suda protocol, also known as low concentration glycerol): Perfusion with glycerol in HBSS at 5%, 10%, and 15% for 20 minutes each at 4°C [19, 20], followed by slow freezing at -20°C (**Fig. S1**). Rewarming and fixation are the same as in condition I.

Condition K (Straight freeze, also known as unprotected cryopreservation): No fixation or cryoprotectant perfusion was performed after the initial PBS wash at 4°C. Specimens were slowly frozen in a Styrofoam container at -20°C within approximately 400 minutes (**Fig. S1**), followed by transfer to -150°C storage. Rewarming and fixation are the same as in condition I.

Condition L (Interleaved equilibration): The methods for this condition were described in more detail previously [22]. Briefly, it consists of perfusion with 59% V3 vitrification solution for 8 minutes, followed by LM5 carrier solution for 3 minutes, followed by 59% V3 for 25 minutes, then vitrification and storage at -150°C. After rewarming, specimens were perfused with 10% dextran for 25 minutes, followed by perfusion fixation with the fixative used in condition B.

The fixation, cryoprotectant loading, and cooling parameters of the twelve preservation conditions are summarized in **Table 1** and schematically represented in **Fig. 1**.

To assess vascular perfusability and cerebral dye distribution under the ASC warm and ASC cryo conditions, methylene blue was perfused at a dilution of 1:10,000 in the respective carrier solution. The excised brains were examined macroscopically for the distribution of staining (Fig. 2F).

**Figure 2.**
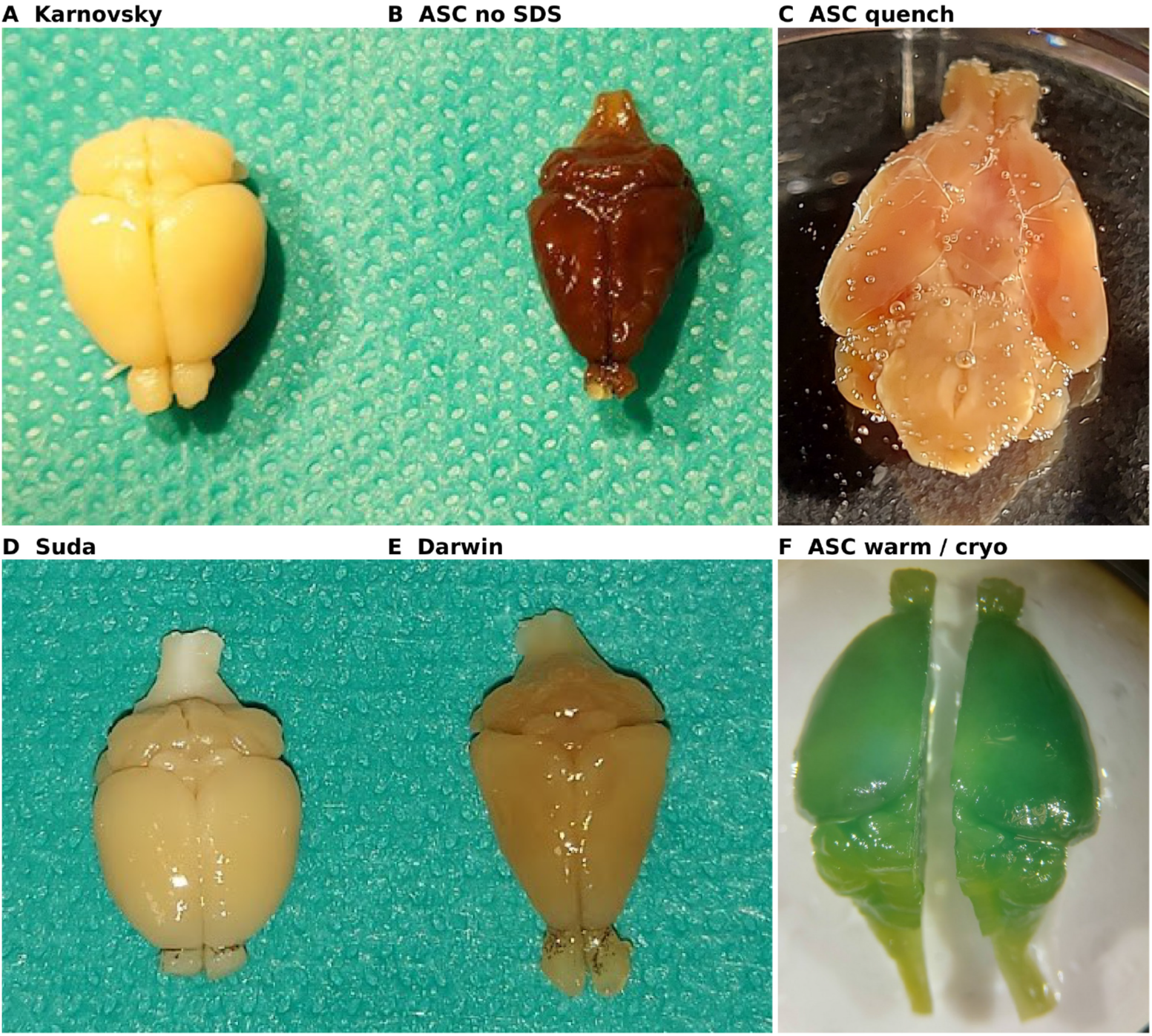
Gross morphology of mouse brains following different preservation protocols. (A) Karnovsky perfusion fixation showing pale yellow discoloration and preserved cortical convexity. (B) ASC without SDS showing dark brown discoloration and near-complete loss of cortical convexity, consistent with severe dehydration. (C) ASC with perfusion quenching, showing isopentane bubbles on the brain surface. (D) Low-concentration glycerol perfusion (Suda protocol). (E) High-concentration glycerol perfusion (Darwin protocol) showing slight brownish discoloration with volume loss. (F) ASC warm (left) versus ASC cryo (right) following methylene blue perfusion to assess vascular patency. Both specimens appear dark green, consistent with intact perfusability and marked cerebral dye uptake due to BBB permeabilization with SDS.

### Freezing experiments to test whether retaining intravascular blood contents affected ice-mediated tearing

To further characterize the effects of freezing on non-perfused aldehyde fixed tissue, additional experiments were performed with mice and rats sacrificed without perfusion via decapitation. In mice, the brain was extracted and immersion fixed in 1% (w/v) paraformaldehyde and 2% (w/v) glutaraldehyde at 4°C for 4 weeks, followed by slow freezing as in condition H to test whether retaining intravascular blood contents affected ice-mediated tearing. In rats, the brain was extracted and immersion fixed with 4% paraformaldehyde alone at 4°C for 1 week. After this, the left hemispheres were stored at 4°C and the right hemispheres subjected to subsequent slow freezing to -20°C as used in condition H. In total, six hemispheres from three rats were used.

### Tissue processing and microscopy

After preservation, samples from folium IV/V of the cerebellar vermis were dissected into approximately 1 mm^3^ specimens. For all preservation conditions, the 1 mm^3^ specimens were immersed in Ito-Karnovsky fixative for 24 hours at 4°C, washed overnight in cacodylate buffer, postfixed with OsO4, dehydrated through a graded ethanol series, and embedded in Epon (Roth, Germany). Semithin sections (1 μm) were stained with toluidine blue and imaged at 4x, 10x, 20x, and 40x magnification on a light microscope.

Ultrathin sections (50 nm, Reichert Ultracut) were subsequently stained with uranyl acetate and lead citrate and viewed with a Zeiss EM 900N at 80 kV, and images were acquired with a side-entry CCD camera (TRS). For each condition, representative images of cerebellar granule cell nuclei, myelin, capillaries, and synapses were acquired.

### Patch-level feature analysis

Patch-level feature analysis was performed using the DINOv2 vision foundation model (dinov2_vitg14_reg, ViT-G/14 architecture with registers) [31]. Each electron micrograph was cropped to remove metadata, resized to 518 × 518 pixels, center-cropped, and normalized (mean = 0.5, SD = 0.2).

The model extracted a 1536-dimensional class token as an image-level representation and 37 × 37 patch tokens as patch-level representations per image. For image-level analysis, class token features from all specimens were embedded in two dimensions using t-distributed stochastic neighbor embedding (t-SNE) with a perplexity of 3 via the python package Scikit-learn. For patch-level visualization, the first three principal components of patch token features were mapped to RGB channels and superimposed on the original micrograph. To quantify ultrastructural similarity between conditions via a patch-level similarity analysis, a vector similarity search was performed using the python package PyNNDescent for fast Approximate Nearest Neighbors [32]. Mean Euclidean distances and cosine similarities of patch-level features to the 10 nearest neighbors from reference conditions were computed and displayed as heatmaps using a jet colormap superimposed on the source images. For the slow-freeze analysis, four independent 4×4 tile grids were stitched (518-pixel tiles with 8-pixel overlap) to assess spatial consistency across a larger field of view.

## Results

### Gross brain examination

The preservation protocols produced marked differences in gross brain morphology (**Fig. 2**). The Karnovsky-fixed brain (condition A) had the pale yellow color typical of picric-acid-containing fixative, a firm texture and a smooth cortical convexity (Fig. 2A). ASC without SDS (condition F) produced a small, dark brown brain with pronounced cortical folding and near-complete loss of convexity, consistent with severe dehydration (Fig. 2B), as described in the original ASC report [25]. The ASC quench specimen (condition E) showed isopentane bubbles on its surface (Fig. 2C). High-concentration glycerol (condition I) produced a shrunken, wrinkled cortex with brownish discoloration (Fig. 2E), whereas low-concentration glycerol (condition J) caused only a slight reduction in brain width (Fig. 2D). ASC warm and ASC cryo brains (conditions C and D) retained normal gross size and shape. Methylene blue perfusion produced evenly distributed dark green staining, consistent with thorough perfusion and cerebral dye uptake across the SDS-permeabilized BBB (Fig. 2F). The SDS perfusion step also produced a milky effluent, suggesting washout of solubilized tissue components.

### Light microscopy

Toluidine blue-stained semithin sections showed recognizable cerebellar architecture across all conditions at 4x, 10x, 20x and 40x magnification (Figs. S2–S4 and **Fig.** 3). Staining intensity varied, with particularly dark staining after ASC without SDS (condition F), consistent with dehydration and tissue compaction. Differences in staining duration may also contribute to this variation.

**Figure 3.**
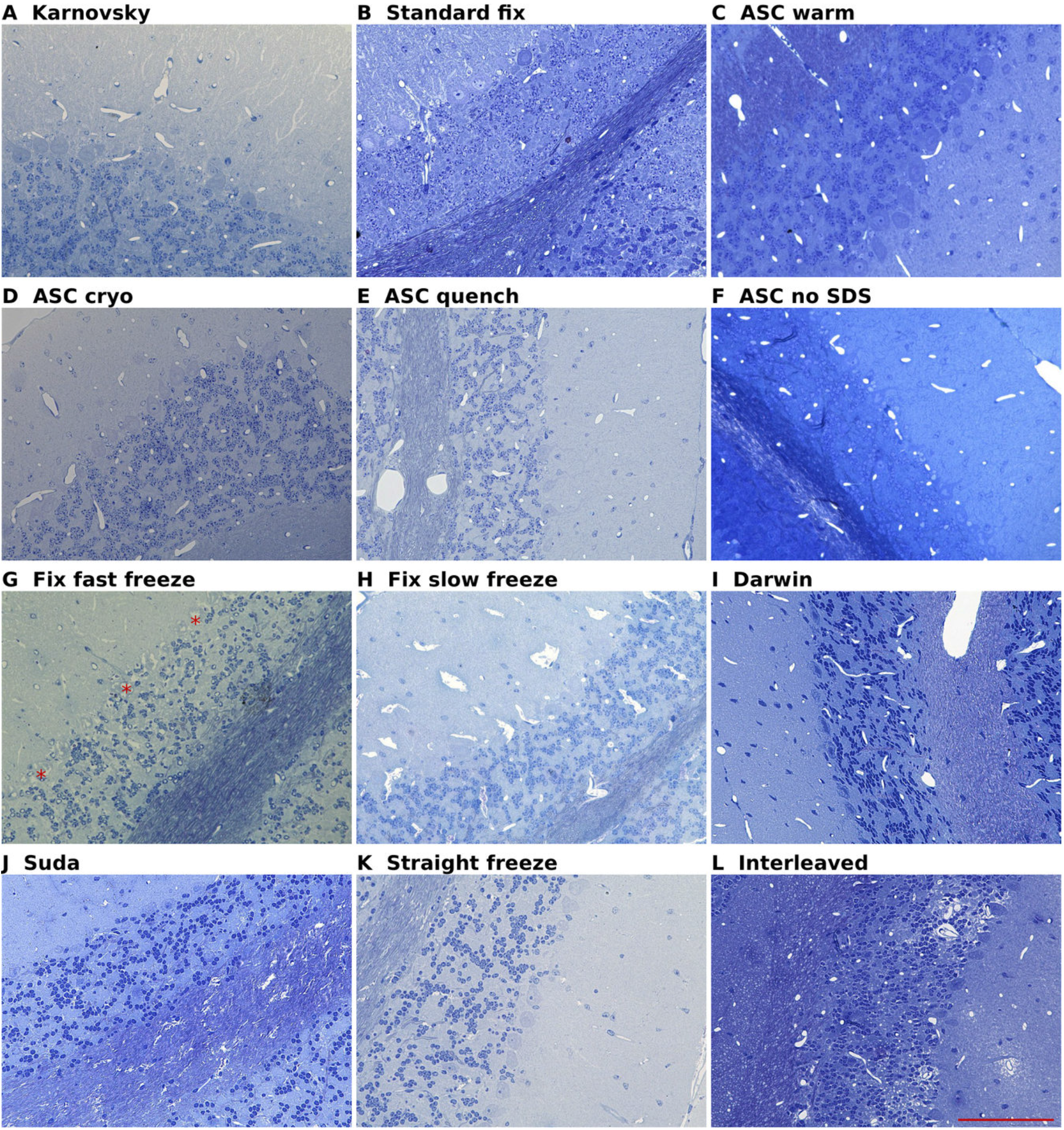
Light microscopy of cerebellar tissue across all twelve preservation conditions at 40x magnification. Toluidine blue-stained semithin sections. (A) Karnovsky fixation, (B) Standard fixation, (C) ASC warm, (D) ASC cryo, (E) ASC quench, (F) ASC without SDS, (G) Fixation followed by fast freezing, (H) Fixation followed by slow freezing, (I) High-concentration glycerol (Darwin), (J) Low-concentration glycerol (Suda), (K) Straight freeze, (L) Interleaved equilibration. Asterisks in G mark poorly delineated Purkinje cell profiles. Scale bar: 100 μm.

Although the layered cerebellar cortex and underlying medulla were clearly seen in all specimens, there were also differences. Aldehyde fixation followed by direct unprotected fast freezing (**Fig. 3G**) presented increased intercellular distances between granule cells and less discernible Purkinje cells, with poorly delineated cell profiles marked by asterisks. Low-concentration glycerol perfusion **(Fig. 3J**) was associated with white matter microtears, and widened perivascular spaces with an amorphous appearance were seen in the protocol using interleaved equilibration with vitrification solution (**Fig. 3L**). Straight freezing retained the gross cerebellar histoarchitecture despite white matter microtears (Fig. 3K).

These observations distinguish the overall retention of cerebellar architecture from protocol-specific changes in cell spacing, perivascular spaces and white matter integrity.

### Electron microscopy

Electron microscopy revealed distinct effects on granule cell nuclei (Fig. 4). Karnovsky and standard fixation, together with SDS-containing ASC (conditions A–E), preserved densely packed cell groups with smooth nuclear contours, dispersed chromatin and intact nuclear envelopes. This appearance persisted after cryogenic cooling in conditions D and **E**. ASC without SDS produced smaller, compact nuclei, consistent with the dehydration seen macroscopically (condition F). Fixation followed by fast freezing produced widely spaced granule cells with small, irregular nuclei, condensed chromatin and intranuclear cavities (condition G). Slow freezing after fixation retained nuclear chromatin and closely packed cells, often around disrupted vessels with widened perivascular spaces (condition H). High-concentration glycerol produced markedly shrunken cells and nuclei that were difficult to distinguish from the condensed cytoplasm, with clefts and cavities separating them from the surrounding parenchyma (condition I). Low-concentration glycerol produced less pronounced shrinkage, irregular nuclear contours and patchy chromatin condensation (condition J). Straight freezing retained identifiable granule cells and nuclei within mildly condensed parenchyma (condition K). Interleaved equilibration preserved recognizable chromatin patterns, with variable nuclear condensation and increased intercellular spaces containing amorphous material (condition L).

**Figure 4.**
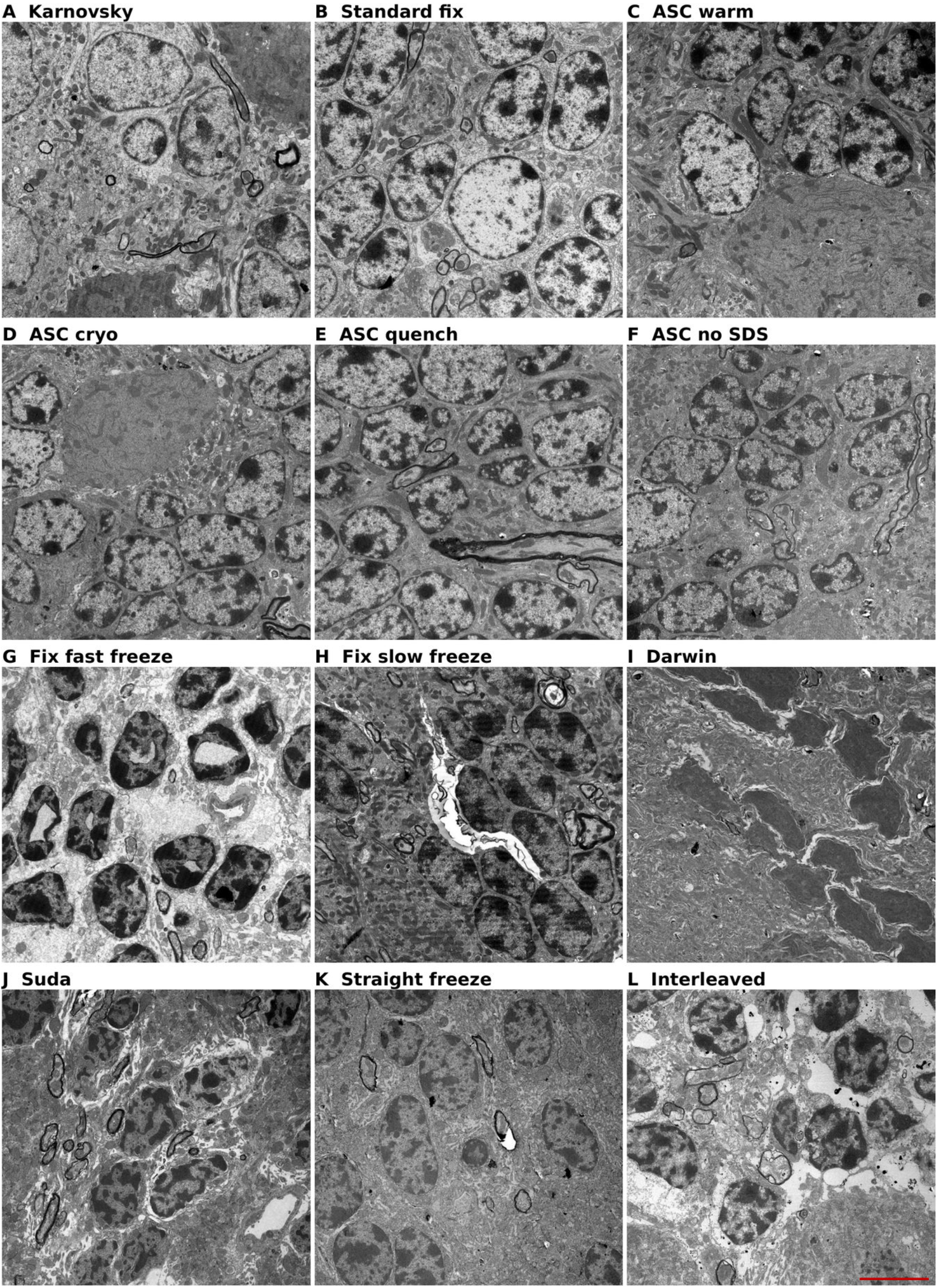
Ultrastructural evaluation of granule cell nuclei in preserved mouse cerebellar tissue. (A) Karnovsky fixation, (B) Standard fixation, (C) ASC warm, (D) ASC cryo, (E) ASC quench, (F) ASC without SDS, (G) Fixation followed by fast freezing, (H) Fixation followed by slow freezing, (I) High-concentration glycerol (Darwin), (J) Low-concentration glycerol (Suda), (K) Straight freeze, (L) Interleaved equilibration. Scale bar: 5 μm.

We next evaluated the ultrastructure of myelin, which we found to be the feature that best distinguished preservation quality across conditions. The standard aldehyde perfusion fixation conditions generally showed well-ordered lamellae, albeit with some areas that were less compact, which is not uncommon in aldehyde-fixed tissue in our experience (**Fig. 5A-E**). ASC without SDS showed mostly intact but shrunken myelin profiles, consistent with tissue compaction from dehydration, and also had some areas of lamellar separation (**Fig. 5F**). Fixation followed by fast freezing showed identifiable myelinated axon profiles but with widespread irregular, delaminated lamellae (**Fig. 5G**), while fixation followed by slow freezing showed well-preserved myelin (**Fig. 5H**). High-concentration glycerol perfusion showed separated myelin profiles with delamination (**Fig. 5I**). Low-concentration glycerol perfusion showed a lower degree of myelin disruption, but with mechanical tearing in the white matter (**Fig. 5J**). Straight freezing without fixation or cryoprotectant showed disrupted myelin architecture with areas containing electron-dense debris (**Fig. 5K**). Finally, the interleaved equilibration method showed well-preserved, identifiable myelinated axon profiles with occasional lamellar disruption (**Fig. 5L**).

**Figure 5.**
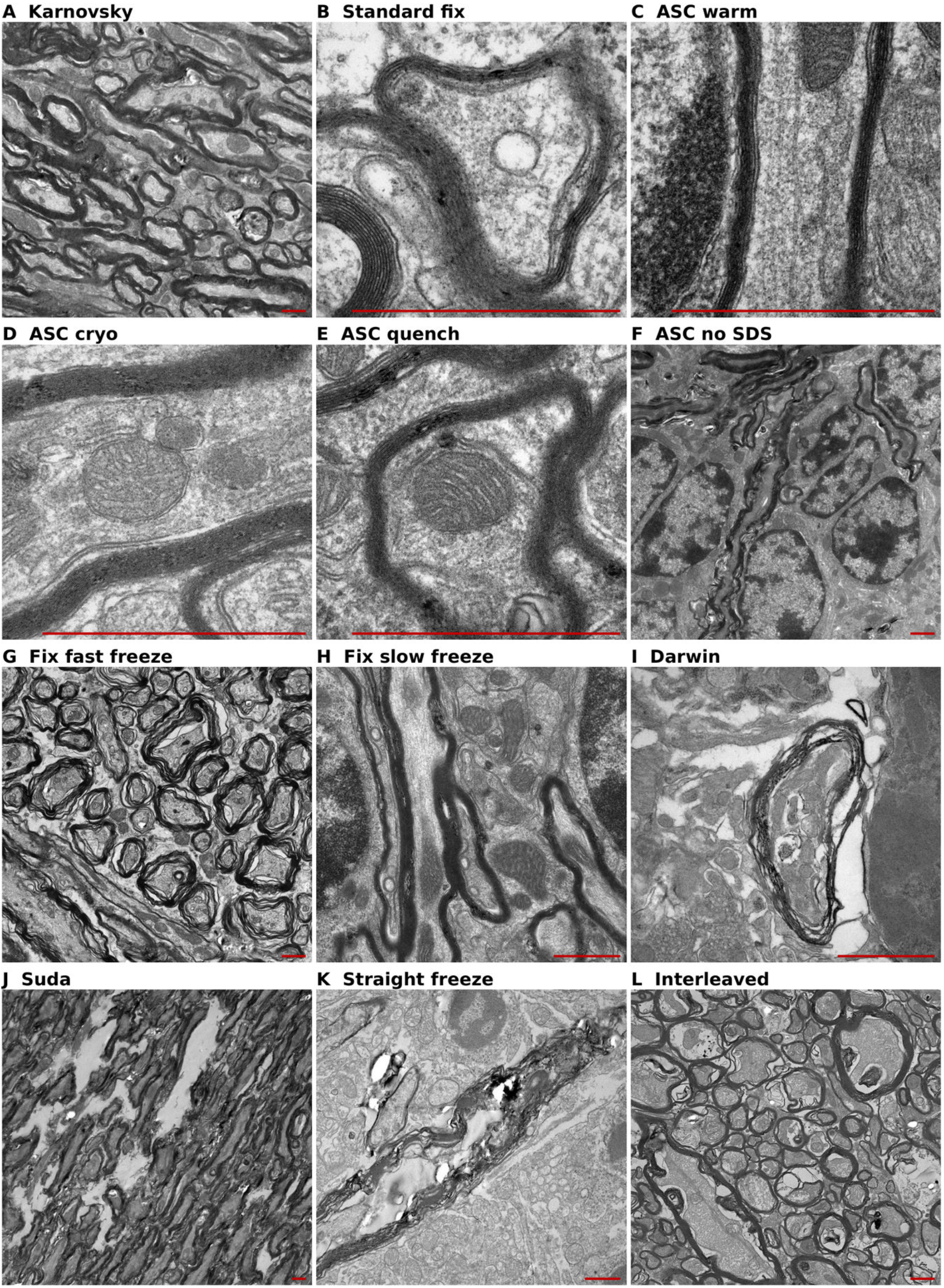
Ultrastructural evaluation of myelin in preserved mouse cerebellar tissue. (A) Karnovsky fixation, (B) Standard fixation, (C) ASC warm, (D) ASC cryo, (E) ASC quench, (F) ASC without SDS, (G) Fixation followed by fast freezing, (H) Fixation followed by slow freezing, (I) High-concentration glycerol (Darwin), (J) Low-concentration glycerol (Suda), (K) Straight freeze, (L) Interleaved equilibration. Scale bars: 1 μm.

Synaptic profiles differed in vesicle definition, membrane continuity and contrast (Fig. 8). Fixation controls and SDS-containing ASC preserved well-defined vesicle clusters and apposed synaptic membranes (conditions A–E). Several alternative protocols showed less distinct membranes and vesicles, with the most pronounced loss of contrast and membrane integrity after straight freezing (condition K). Interleaved equilibration retained recognizable synaptic profiles and substantial local organization (condition L).

### Capillaries

Because vascular access is central to cryoprotectant delivery, we next assessed capillary ultrastructure. The standard aldehyde-fixed conditions, as well as ASC without SDS, showed capillaries with intact endothelial lining and sharply-defined lumens (**Fig. 6A-F**). Fixation followed by fast freezing retained open capillary lumens (**Fig. 6G**), while fixation followed by slow freezing showed perivascular microtears and endothelial disruption, with additional membrane-bound profiles and electron-dense material within the lumen (**Fig. 6H, S7**). Glycerol perfusion conditions showed relatively preserved capillary profiles, though with shrunken areas in the surrounding tissue (**Fig. 6I**, **Fig. 6J**). Straight freezing without fixation or cryoprotectants showed debris-like structures within the capillary lumen, with the vessel profile marked by an asterisk (**Fig. 6K**). Interleaved equilibration with vitrification solution showed well-defined vessel lumens but with widened perivascular space (**Fig. 6L**).

**Figure 6.**
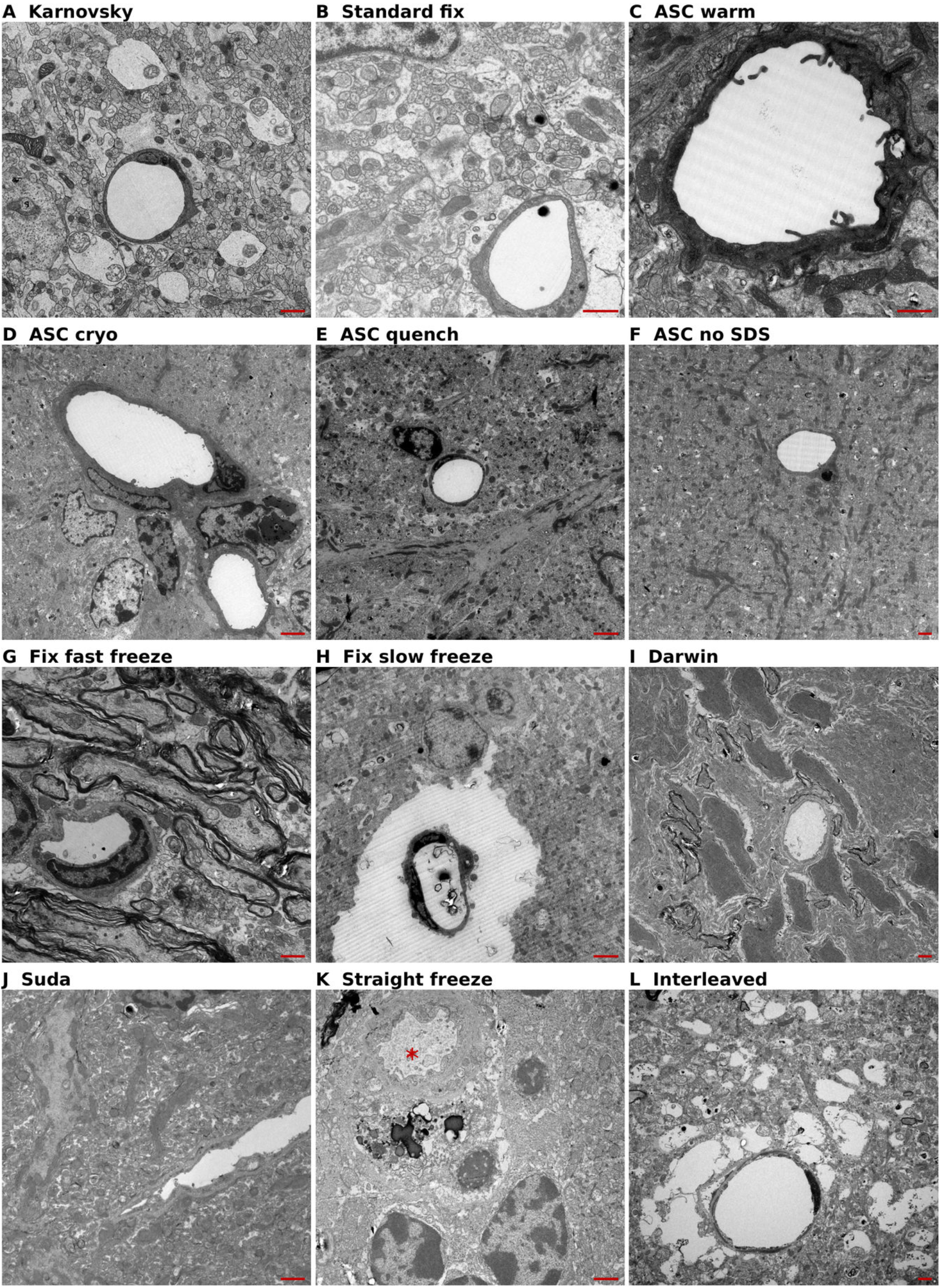
Ultrastructural evaluation of capillaries in preserved mouse cerebellar tissue. (A) Karnovsky fixation, (B) Standard fixation, (C) ASC warm, (D) ASC cryo, (E) ASC quench, (F) ASC without SDS, (G) Fixation followed by fast freezing, (H) Fixation followed by slow freezing, (I) High-concentration glycerol (Darwin), (J) Low-concentration glycerol (Suda), (K) Straight freeze, (L) Interleaved equilibration. The asterisk marks the vessel profile in K. Scale bars: 1 μm.

Additional images after interleaved equilibration showed widened perivascular spaces in multiple cerebellar vessels, consistent with edema arising during CPA loading or unloading (Fig. 7A, C). Some vessels also showed focal endothelial disruption (**Fig. 7E**). The surrounding parenchyma retained recognizable neuropil and myelinated axon profiles (**Fig. 7F**), placing the dominant injury at the neurovascular interface.

**Figure 7.**
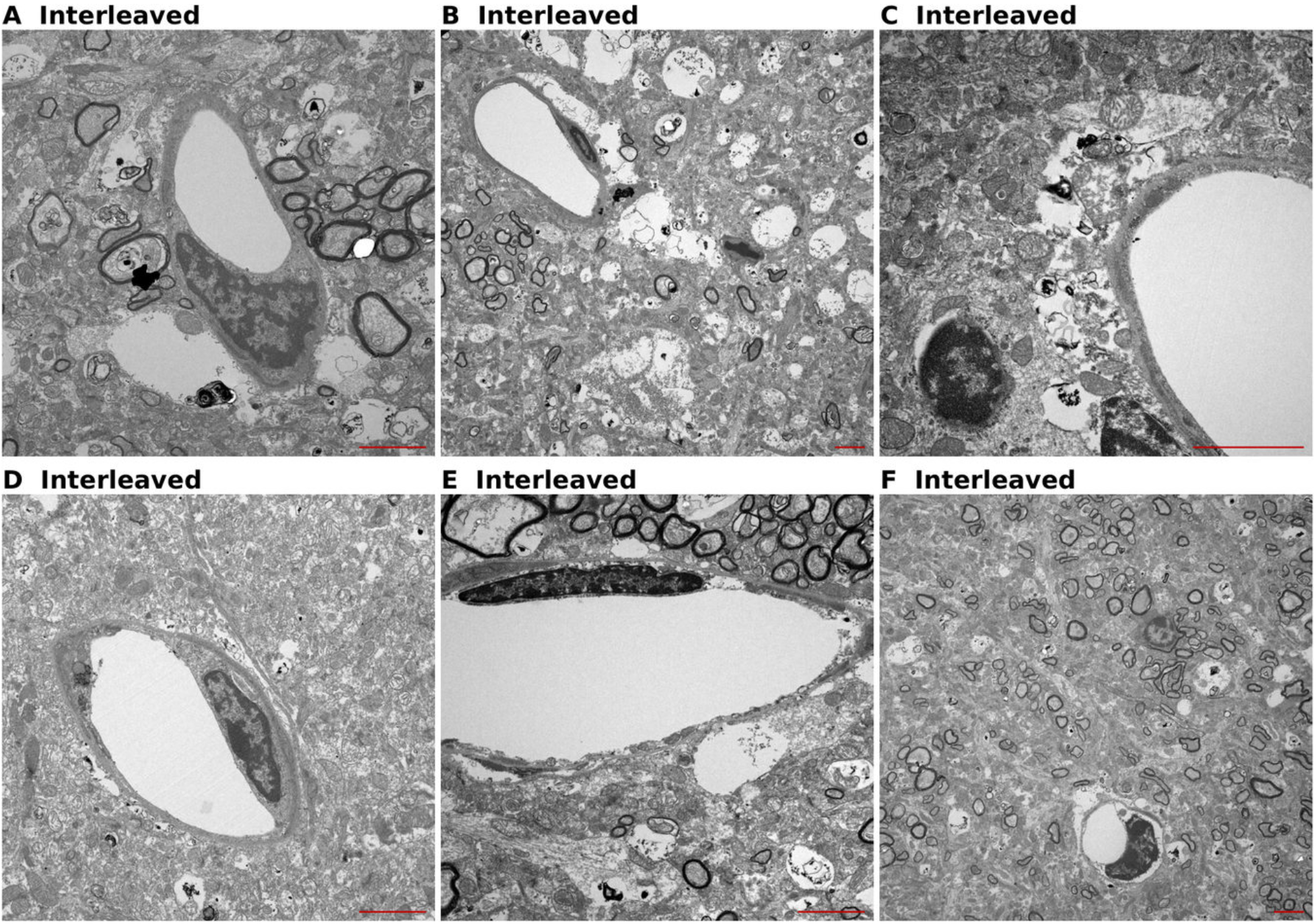
Ultrastructural evaluation of cerebellar capillaries following interleaved equilibration with vitrification solution. Multiple vessel profiles showing widened perivascular spaces consistent with edema (A, C) and focal endothelial disruption (E). The surrounding parenchyma retains identifiable neuropil texture and myelinated axon profiles (F). Scale bars: 2 μm.

**Figure 8.**
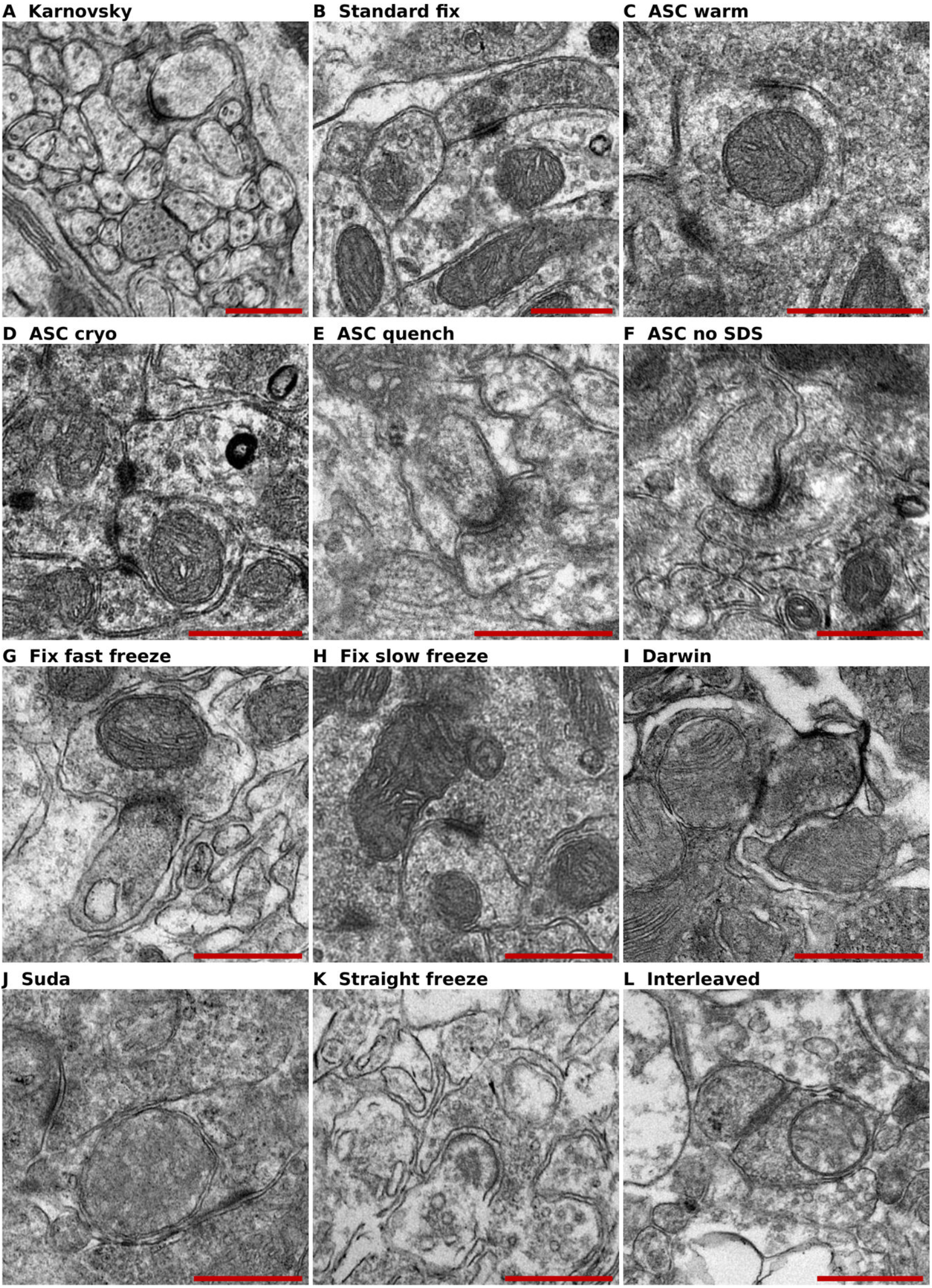
Ultrastructural evaluation of synapses in preserved mouse cerebellar tissue. (A) Karnovsky fixation, (B) Standard fixation, (C) ASC warm, (D) ASC cryo, (E) ASC quench, (F) ASC without SDS, (G) Fixation followed by fast freezing, (H) Fixation followed by slow freezing, (I) High-concentration glycerol (Darwin), (J) Low-concentration glycerol (Suda), (K) Straight freeze, (L) Interleaved equilibration. Scale bars: 500 nm.

We further characterized the dehydration artifact observed when tissue was preserved via ASC without SDS by examining forebrain tissue from the same specimen (**Fig. S5**). We found that near the glia limitans, the tissue appeared relatively preserved, with more easily identifiable cellular structures (**Fig. S5A**). This superficial-to-deep gradient points to a mismatch between water and cryoprotectant transport at the brain surface. Direct exposure of the superficial tissue to cryoprotectant provides one explanation for the relatively preserved surface layer.

### Aldehyde fixation followed by unprotected freezing

To further characterize the effects of freezing on aldehyde-fixed tissue, we performed follow-up experiments varying fixation and freezing conditions. We evaluated additional ultrastructural images to corroborate that glutaraldehyde fixation followed by fast freezing via liquid nitrogen immersion leads to intranuclear cavities (**Fig. S6**). We also corroborated that when the tissue was fixed via glutaraldehyde and then frozen slowly, the dominant artifact was perivascular microtearing (**Fig. S7**). These tears appeared as linear clefts radiating from or adjacent to capillary profiles. In perfused tissue, the vascular lumen is filled with aqueous solution rather than blood, which may provide a low-barrier pathway for ice propagation [33], leading to perivascular mechanical tears. Aside from this artifact, parenchymal areas generally had recognizable structures, suggesting that ice damage in slowly frozen fixed brain tissue is predominantly found in perivascular spaces.

To test whether keeping blood contents in the vasculature during slow freezing might reduce mechanical tearing, we performed a follow-up experiment using immersion fixation with glutaraldehyde rather than perfusion fixation prior to slow freezing. Using light microscopy, we found that slow freezing still produced tissue microtears, suggesting that retaining intravascular contents does not prevent ice-mediated mechanical disruption (**Fig. S8**).

We next asked whether the choice of fixative modulates the susceptibility to freeze-induced damage. We found that the immersion fixation of rat cerebellum with PFA alone (without glutaraldehyde) yielded structural preservation comparable to perfusion fixation conditions when the tissue was stored at 4°C prior to embedding (**Fig. 9A**, **Fig. 9C**). However, subsequent slow freezing of PFA-fixed tissue produced a severe Swiss-cheese pattern of vacuolation that affected all cortical layers and was easily visible even on light microscopy (**Fig. 9B**). This result was replicated three times independently. The preservation outcome was markedly worse than what was observed when glutaraldehyde-containing fixative was used prior to slow freezing, in which case the damage was largely confined to perivascular zones. This may be because glutaraldehyde fixation creates a more rigid gel-like structure than PFA fixation, thus making it more resistant to the mechanical forces generated by extracellular ice crystal growth during slow cooling and decreasing tissue displacement [3].

**Figure 9.**
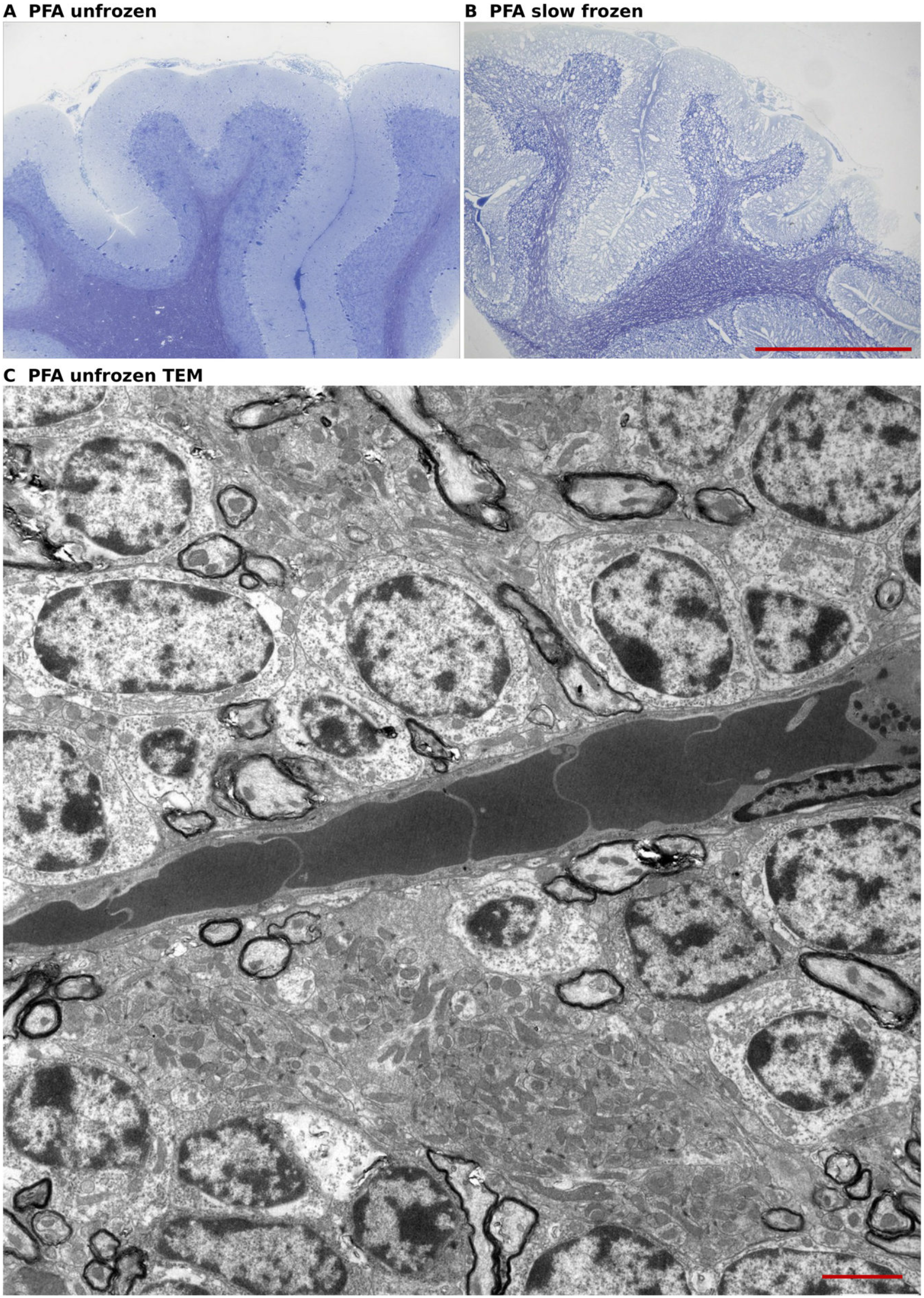
Effect of slow freezing on PFA-fixed rat cerebellar tissue. (A) Immersion fixation with PFA alone showing preserved tissue-level architecture. (B) PFA fixation followed by slow freezing showing severe vacuolation in a Swiss-cheese pattern across all cortical layers. (C) Electron microscopy of PFA-fixed tissue prior to freezing, with erythrocytes visible in the vasculature due to the lack of perfusion. Scale bars: 500 μm (A, B), 2.5 μm (C).

### Patch-level feature analysis

To complement the qualitative ultrastructural assessment, we performed image analysis using the DINOv2 vision foundation model (Fig. 10). Principal component visualization of patch-level features revealed that the model captured biologically meaningful texture variation, with distinct colorization of neuropil, myelin, and nuclear compartments without domain-specific training (**Fig. 10A**). A t-SNE embedding of image-level features showed that image magnification is the dominant source of variance, although there was also some clustering by the preservation condition (**Fig. 10B**). Patch-level distance heatmaps relative to non-vitrified ASC showed low distances for other standard methods of perfusion fixation, and markedly elevated distances for fast freezing, where the highest distances localized to intranuclear cavities and disrupted myelin. ASC without SDS showed moderately elevated distances that were consistent with dehydration (**Fig. 10C**). Cosine similarity analysis of glutaraldehyde fixation followed by slow freezing relative to Karnovsky fixation highlighted perivascular tears and, less consistently, regions adjacent to myelin sheaths. The remaining parenchyma resembled the fixation-only control condition (**Fig. 10D**).

**Figure 10.**
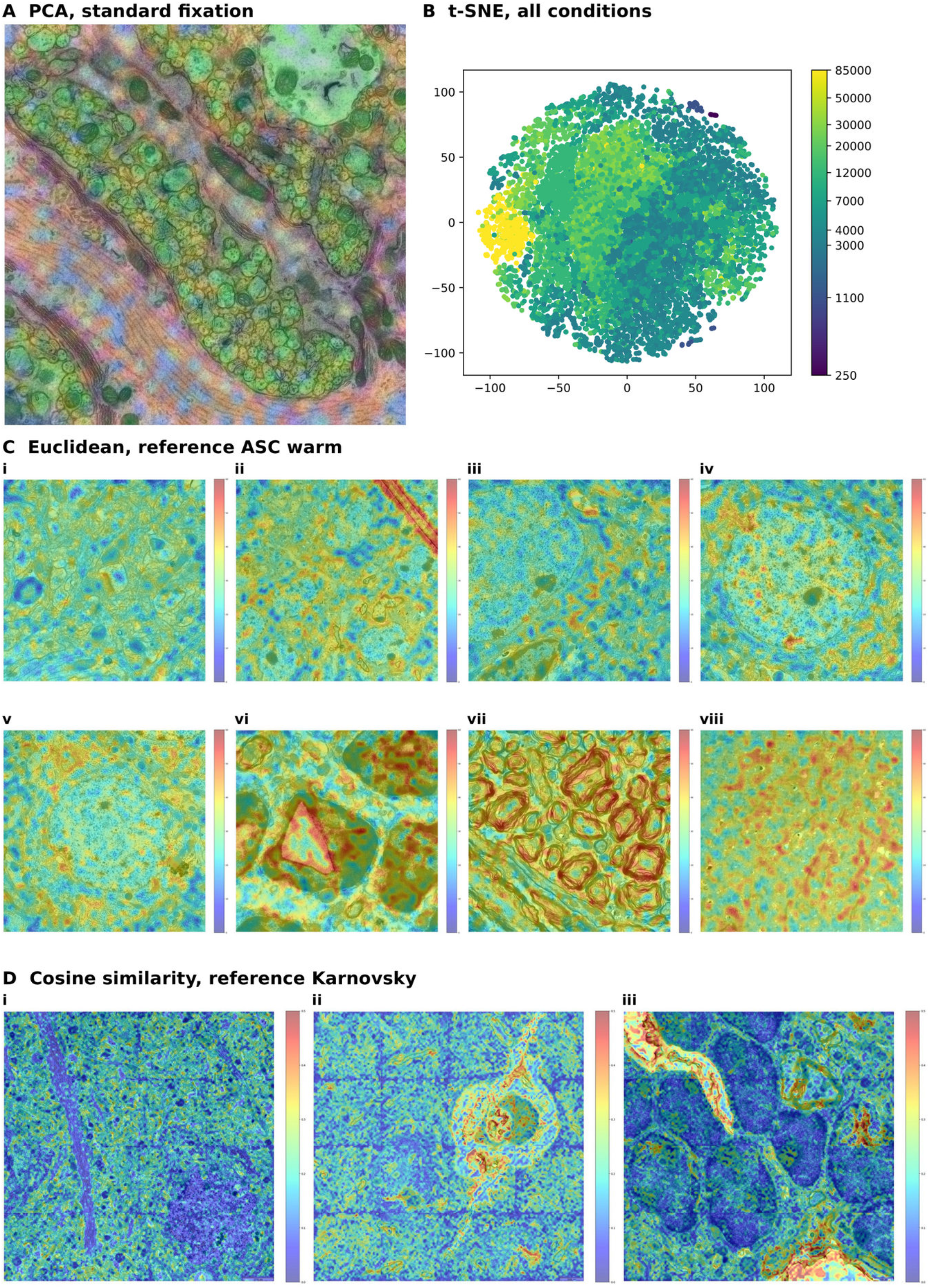
Patch-level feature analysis of ultrastructural preservation quality. (A) Principal component visualization of patch-level features from the DINOv2 vision foundation model, with the first three components mapped to RGB channels and superimposed on the original micrograph from condition B. (B) t-SNE embedding of image-level features from all specimens, with viridis colormap indicating magnification. (C) Mean Euclidean distances of patch-level features to 10 nearest neighbors from condition C (ASC warm). Image-specimen pairs: i, condition B; ii, condition D (ASC cryo, with diamond knife cutting artifact); iii, condition E (ASC quench); iv, condition A; v, condition B (fixation controls); vi-vii, condition G (fast freezing, with intranuclear cavities and disrupted myelin); viii, condition F (ASC without SDS). Jet colormap scaled 0-60. (D) Mean cosine similarity of patch-level features to 10 nearest neighbors from condition A (Karnovsky fixation), with 4×4 tile grids stitched to assess spatial consistency. Image-specimen pairs: i, condition A (self-comparison); ii-iii, condition H (slow freezing). Differences in cosine similarity localize to perivascular tears and myelin sheaths. Jet colormap scaled 0-0.5.

### Summary of methods differences

Table 2 summarizes the light microscopic and ultrastructural findings across the twelve preservation conditions.

**Table 2.** Summary of light microscopy and ultrastructural findings across the twelve preservation conditions.

| Condition | Light microscopy (semithin sections) | Ultrastructure (TEM) |
| --- | --- | --- |
| A | Preserved folial contour and lamination. | Preserved nuclei, myelin, microvasculature, and synaptic neuropil. |
| B | Overall preserved lamination; occasional hyperchromatic (dark) neurons. | Comparable to A. |
| C | Comparable to A and B. | Comparable to A and B. |
| D | Comparable to A and B. | Comparable to A and B. |
| E | Comparable to A and B. | Comparable to A and B. |
| F | Hyperstaining with maintained laminar organization. | Increased electron density and heterogeneity in neuropil; sharp dehydration/staining gradient at the glia limitans in samples from the forebrain. |
| G | Purkinje cell layer disruption and geometric intranuclear clearing. | Intranuclear angular void areas with associated neuropil disruption and vacuolization. |
| H | Perivascular microtears. | Perivascular microtears and varying endothelial disruption. Other domains comparable to A and B. |
| I | Nuclear shrinkage and hyperchromasia with perinuclear halo-like clearing. | Severe nuclear shrinkage and osmophilia with perinuclear halo-like clearing; variable myelin disruption. Identification of cellular compartments and organelles is difficult. |
| J | White matter microtears; mild chromatin clumping. | Myelin disruption. Less severe dehydration and osmophilia than condition I. |
| K | White matter microtearing. Otherwise comparable to A and B. | Filling of capillaries with electron-dense material, disruption of membrane and myelin integrity. |
| L | Perivascular edema. | Perivascular edema with focal endothelial disruption. |

**Abbreviation:** TEM, transmission electron microscopy.

## Discussion

SDS-containing ASC preserved murine cerebellar ultrastructure at a level comparable to standard aldehyde perfusion fixation. Nuclear morphology, myelin periodicity and neuropil texture remained well preserved after cryoprotectant loading alone (condition C) and after subsequent cryogenic cooling (conditions D and E). The alternative protocols produced distinct patterns of dehydration, edema and ice-mediated disruption, identifying the principal biophysical challenges for brain preservation.

### Replication and extension of aldehyde-stabilized cryopreservation

We confirm two key findings from the original ASC report [25]. First, that perfusion fixation followed by SDS-mediated BBB permeabilization and graded ethylene glycol loading preserves ultrastructure at a level comparable to standard aldehyde perfusion fixation alone. Second, that omitting the SDS step results in significant parenchymal dehydration, attributed to inadequate cryoprotectant penetration across the intact BBB. This was evident both macroscopically as near-complete loss of cortical convexity and microscopically as increased tissue compaction. In forebrain tissue from the ASC-without-SDS specimen, we observed a sharp superficial-to-deep dehydration gradient near the glia limitans, indicating a regional mismatch between water and cryoprotectant transport at the brain surface, with direct cryoprotectant exposure providing a route to the superficial tissue.

ASC combines the structural fidelity of aldehyde fixation with access to long-term cryogenic storage. Its value for brain banking lies in stabilizing the tissue before cooling and suppressing the progressive molecular degradation that occurs during storage in liquid fixative. Antigenicity in aldehyde-fixed tissue can decline over days to years, depending on the antigen [4]. The choice of fixation and storage conditions therefore depends on the molecular readouts required alongside ultrastructure.

The fixative also determines which molecular assays remain accessible. Glutaraldehyde provides strong ultrastructural stabilization, while its protein crosslinks can impair antibody recognition. Using antibodies raised against fixed antigens offers one way to combine this structural advantage with immunocytochemical analysis [5].

The milky effluent during SDS perfusion suggests washout of solubilized tissue components. SDS can remove substantial cellular material at higher concentrations and longer exposure times, as illustrated by rat kidney decellularization with 0.66% SDS for 2 hours [34]. ASC uses 0.01% SDS for 20 minutes after fixation. Defining the lipid loss under these conditions will clarify its compatibility with lipidomics and other assays sensitive to membrane composition. Effective BBB permeabilization is central to the structural success of ASC, making selective control of permeability an important target for further optimization.

### Unfixed cryoprotectant perfusion protocols

The glycerol-based protocols (conditions I and J) were associated with nuclear shrinkage and hyperchromasia, consistent with osmotic dehydration. The high-concentration glycerol protocol (condition I, Darwin protocol) produced particularly severe nuclear shrinkage, making identification of cellular compartments difficult. This was somewhat surprising, as the original report identified generally normal ultrastructure aside from ice crystal damage in select areas [28]. The low-concentration glycerol protocol (condition J, Suda protocol) showed less severe but still notable dehydration and myelin disruption. These findings suggest that glycerol loading and unloading without prior fixation imposes osmotic stress on the tissue, and that the unfixed brain parenchyma is particularly vulnerable to the volume shifts associated with cryoprotectant equilibration.

Natural freeze tolerance couples endogenous cryoprotectant accumulation to metabolic acclimation [12]. Maintaining cellular metabolism during equilibration allows volume regulation and adaptation to changes in water content, ionic strength and osmolarity. The associated metabolic, gene-expression and protein responses offer targets for improving mammalian tissue tolerance to CPA loading.

Interleaved equilibration with vitrification solution (condition L) preserved identifiable myelinated axons, chromatin patterns and synaptic profiles, supporting its value for applications requiring unfixed tissue. Its principal artifacts were perivascular edema and focal endothelial disruption. Optimizing CPA loading and unloading at the BBB should improve structural preservation while retaining the functional and molecular advantages of avoiding aldehyde crosslinking.

The BBB separates cerebral fluid from blood plasma through the luminal and abluminal endothelial membranes and the tight junctions between endothelial cells [35, 36]. The relative contributions of transcellular and paracellular routes to water transport remain unresolved. Beyond the endothelium, water crosses the perivascular compartment and aquaporin-4-rich astrocyte endfeet, which can limit water exchange in knockout models [37]. Water and CPA permeability differ substantially: a water exchange fraction of 0.84 per perfusion pass has been reported in the human brain at 36°C [38], while permeability in bovine pulmonary artery endothelial cells at 4°C follows the order water > formamide > DMSO > ethylene glycol > glycerol [39]. The broad parenchymal dehydration observed here points to a mismatch at a shared transport barrier. A limitation confined to individual cell membranes would be expected to produce more differential shrinkage among cell populations. The preserved superficial layer overlying a dehydrated interior near the glia limitans also identifies a barrier to water and CPA equilibration at the brain surface **(Fig. S5A**). Dehydration increased with glycerol concentration, persisted despite aldehyde fixation and room-temperature CPA loading without SDS, and disappeared when the BBB was permeabilized with SDS. Interleaving hypotonic carrier reversed the volume artifact toward edema and was accompanied by endothelial discontinuities (**Fig.** 7). Together, these observations locate the major transport limitation and mechanical vulnerability at the neurovascular interface. Loading temperature and CPA composition offer direct means of adjusting the balance between water flux and CPA entry.

The luminal and abluminal endothelial membranes impose sequential barriers to transcellular CPA transport. Their lipid composition differs: cultured brain capillary endothelial cells show apical enrichment of phosphatidylcholine and basolateral enrichment of sphingomyelin and glucosylceramide [40]. This polarity makes the two membrane domains distinct targets for improving CPA permeability.

CPA composition also changes membrane behavior. Molecular simulations show that DMSO increases membrane fluidity and can induce transient water pores at higher concentrations [41]. These effects provide a mechanistic link between CPA entry, water flux and the vascular injury seen during interleaved equilibration.

### Structural restitution after straight freezing

Straight freezing retained more recognizable architecture after thawing than the extent of ice formation might suggest. Structural restitution offers a mechanistic explanation: cells distorted and compressed by extracellular ice can rehydrate and regain much of their morphology upon thawing [42, 43]. During slow freezing, extracellular ice draws water from cells and displaces tissue. Melting reverses the osmotic gradient and permits re-expansion [2]. Consistent with this process, frozen nerve tissue has shown better apparent preservation after thawing and fixation than after freeze substitution [44].

Structural restitution explains the recovery of recognizable profiles, while membrane disruption and uncertain local displacements limit their use for nanoscale reconstruction. The extensive vacuolation after slow freezing of PFA-fixed tissue illustrates the forces generated by ice growth. In unfixed tissue, re-expansion during thawing can conceal part of this displacement. Straight freezing therefore provides a useful morphological baseline, with cryoprotection required when faithful preservation of fine spatial relationships is the goal.

While comparative biology supports the plausibility of brain tissue surviving controlled partial freezing at shallow subzero temperatures with bounded ice fractions, the unadapted mammalian nervous system is thus not inherently freeze-tolerant. Our results will not apply to cryoprotectant-based methods presently applied in the cryonic preservation of decedents, due to the post-mortem breakdown of the blood-brain barrier [45, 46]. However, condition K hints at the likely ultrastructural state in cases of unprotected freezing.

### Implications for brain banking

Standard aldehyde perfusion fixation remains the reference for morphological studies and connectomics. For long-term storage of structurally stabilized tissue, SDS-containing ASC followed by cryogenic cooling (conditions D and E) provides the strongest overall preservation among the methods tested.

Interleaved equilibration offers the most promising route for applications requiring unfixed tissue, including electrophysiological studies after recovery, with vascular injury as the principal target for improvement. When detergents are incompatible with the intended assays, CPA delivery by surface diffusion may support preservation of sufficiently small or accessible fixed specimens. More selective endothelial permeabilization, or modulation of water transport through aquaporin-4, could further improve equilibration by reducing the mismatch between water flux and CPA entry.

### Limitations

This exploratory comparison used one mouse per condition and a qualitative ultrastructural assessment, complemented by patch-level image analysis. It identifies characteristic preservation patterns and mechanisms but does not estimate biological variability. Species and scale also differ from earlier studies, which examined ASC in a porcine [25], the Suda protocol in a feline [19], and the Darwin protocol in a canine model [28]. Post-thaw fixation varied: conditions I–K received PFA immersion fixation and condition L received PFA/glutaraldehyde perfusion fixation. All samples subsequently underwent the same Ito-Karnovsky postfixation, providing a common microscopy workflow while retaining the influence of their earlier fixation history. Analysis focused on cerebellar tissue and two-dimensional sections. Extension to other brain regions, volumetric connectome tracing and direct molecular measurements will establish how broadly these preservation patterns apply.

## Conclusion

This study provides the first independent replication of ASC and a comparative account of brain preservation from gross anatomy to ultrastructure. Among the cryogenic methods tested, SDS-containing ASC best preserved nuclear morphology, myelin periodicity and neuropil texture. The alternative protocols reveal how ice growth and mismatched water and CPA transport reshape tissue, with the neurovascular interface emerging as a central target for improvement. These findings guide the choice and development of preservation methods for brain banking and neuroscience.

## Abbreviations

**ASC**: Aldehyde-stabilized cryopreservation; **BBB**: Blood-brain barrier; **CPA**: Cryoprotective agent; DMSO: Dimethyl sulfoxide; **EG**: Ethylene glycol; **GA**: Glutaraldehyde; **HBSS**: Hanks’ Balanced Salt Solution; **PBS**: Phosphate-buffered saline; **PFA**: Paraformaldehyde; **SDS**: Sodium dodecyl sulfate; **TEM**: Transmission electron microscopy; **t-SNE**: t-distributed stochastic neighbor embedding.

## Author contributions

A.G., C.F.K., F.P., and J.W. conceptualized the study. A.G. performed laboratory experiments, patch-level image analysis and prepared the figures. A.G. and A.T.M. performed data analysis and wrote the initial draft of the manuscript. All authors reviewed the manuscript and approved the final manuscript.

## Acknowledgements

We thank Andrea Eichhorn, Elke Kretzschmar, and Holger Meixner for technical help. We are grateful to Johanna Habermeyer and Stephan von Hörsten for technical advice and brain tissue sharing access.

## Funding

German Society of Cryobanks (A.G., J.W.)

German Research Foundation grants SFB 1483, FOR 5534 (A.G.)

Interdisciplinary Center for Clinical Research Erlangen project J111 (A.G.)

German Research Foundation grant 460333672 CRC1540 EBM (project A02 - F.P.).

## Data availability

Code used for data analysis is available at DOI 10.5281/zenodo.22131634. Electron microscopy data is available at DOI 10.5281/zenodo.22131046.

## Declaration of Generative AI and AI-Assisted Technologies

During the preparation of this manuscript, the authors used Claude (Anthropic) and ChatGPT (OpenAI) to improve the manuscript’s language. All AI tool-assisted content was reviewed and edited by the authors, who take full responsibility for the final publication.

## Supplementary Material

### Supplementary Figures

**Supplementary Figure S1.**
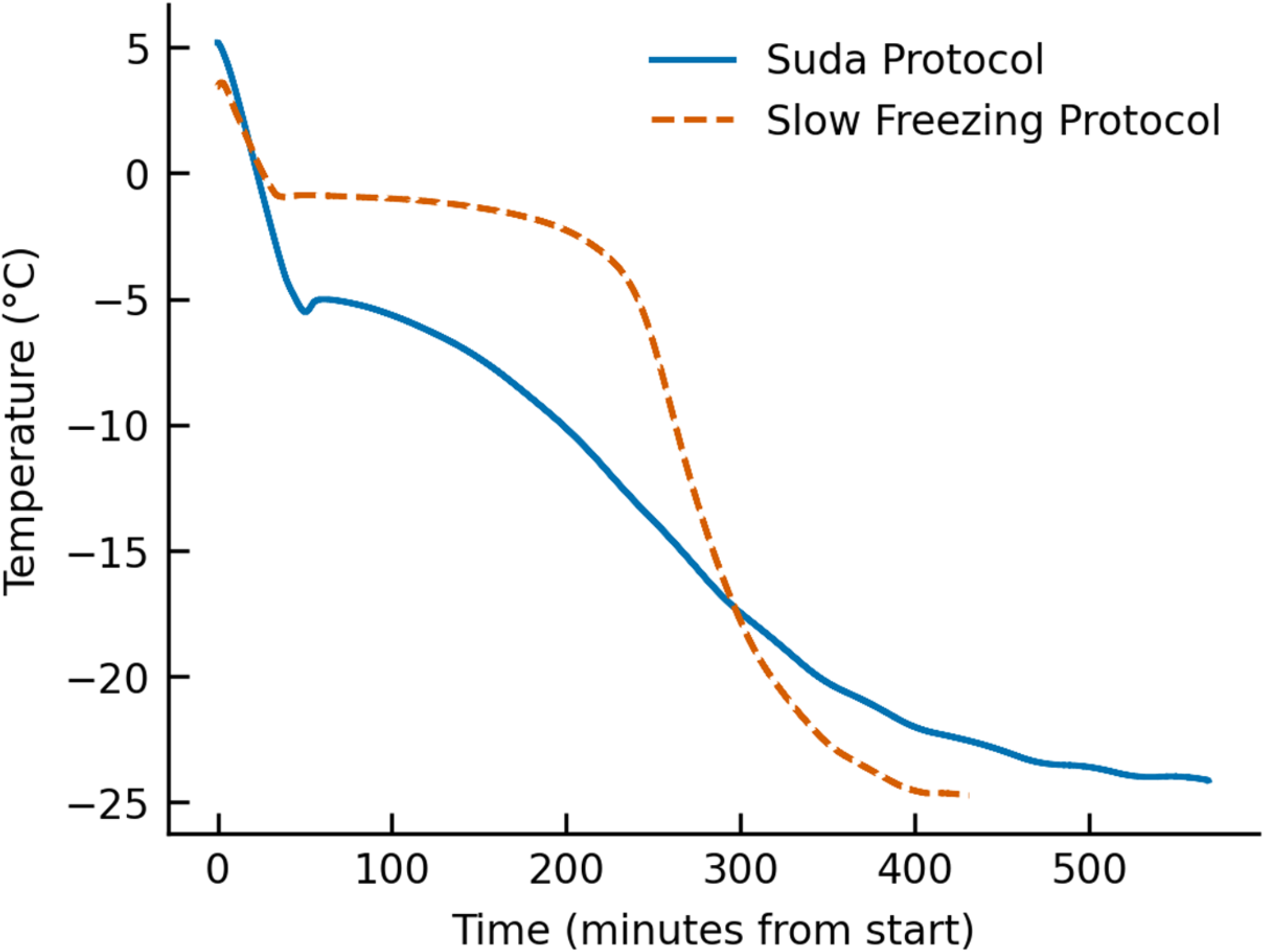
Thermal profiles during slow cooling to -20°C for the Suda protocol (blue) and the slow freezing method used in both Condition H and the straight freeze protocol (orange).

**Supplementary Figure S2.**
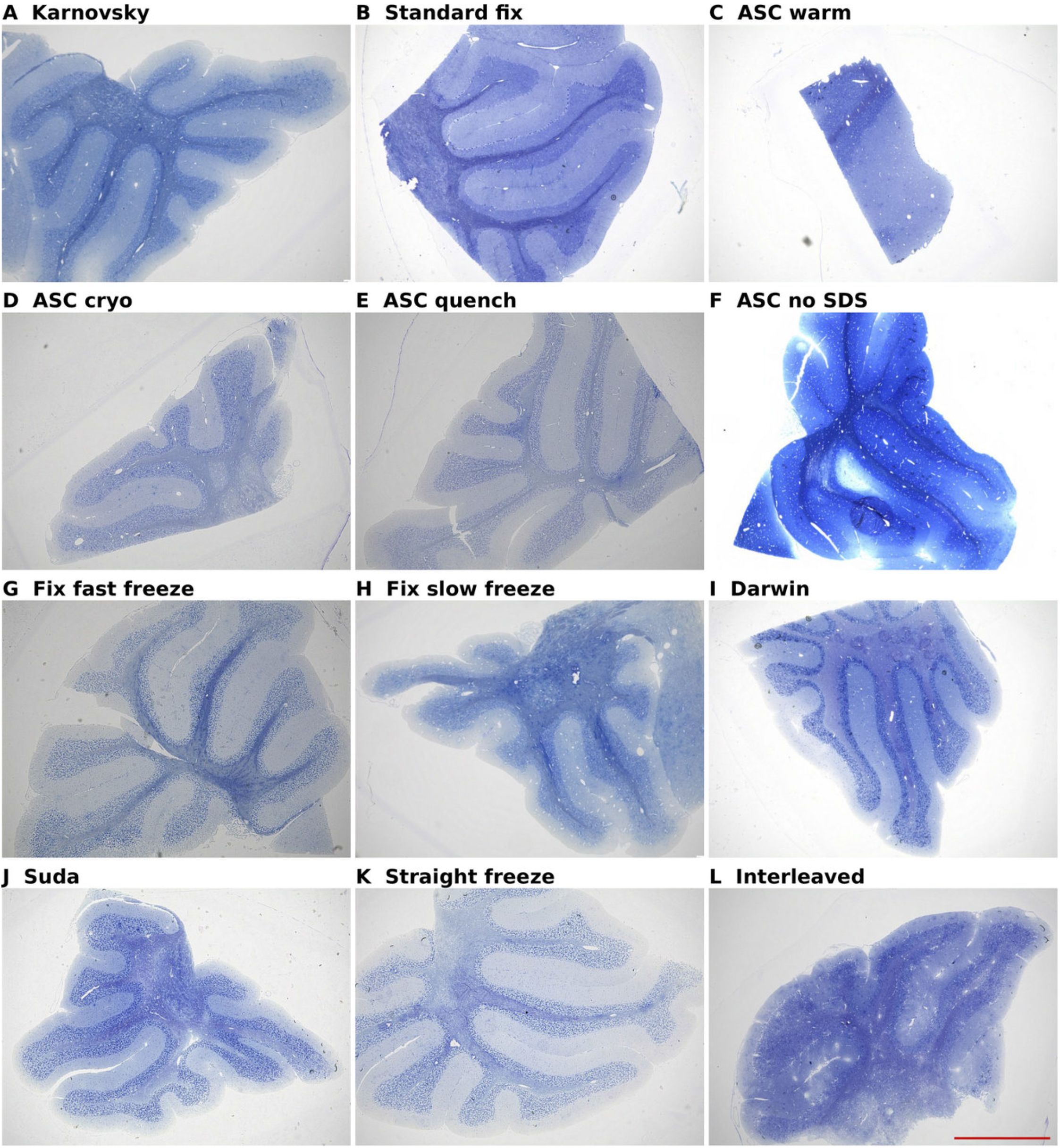
Low-magnification light microscopy of cerebellar tissue across all twelve preservation conditions. Toluidine blue-stained semithin sections at 4x magnification. (A) Karnovsky fixation, (B) Standard fixation, (C) ASC warm, (D) ASC cryo, (E) ASC quench, (F) ASC without SDS, (G) Fixation followed by fast freezing, (H) Fixation followed by slow freezing, (I) High-concentration glycerol (Darwin), (J) Low-concentration glycerol (Suda), (K) Straight freeze, (L) Interleaved equilibration. Scale bar: 1000 μm.

**Supplementary Figure S3.**
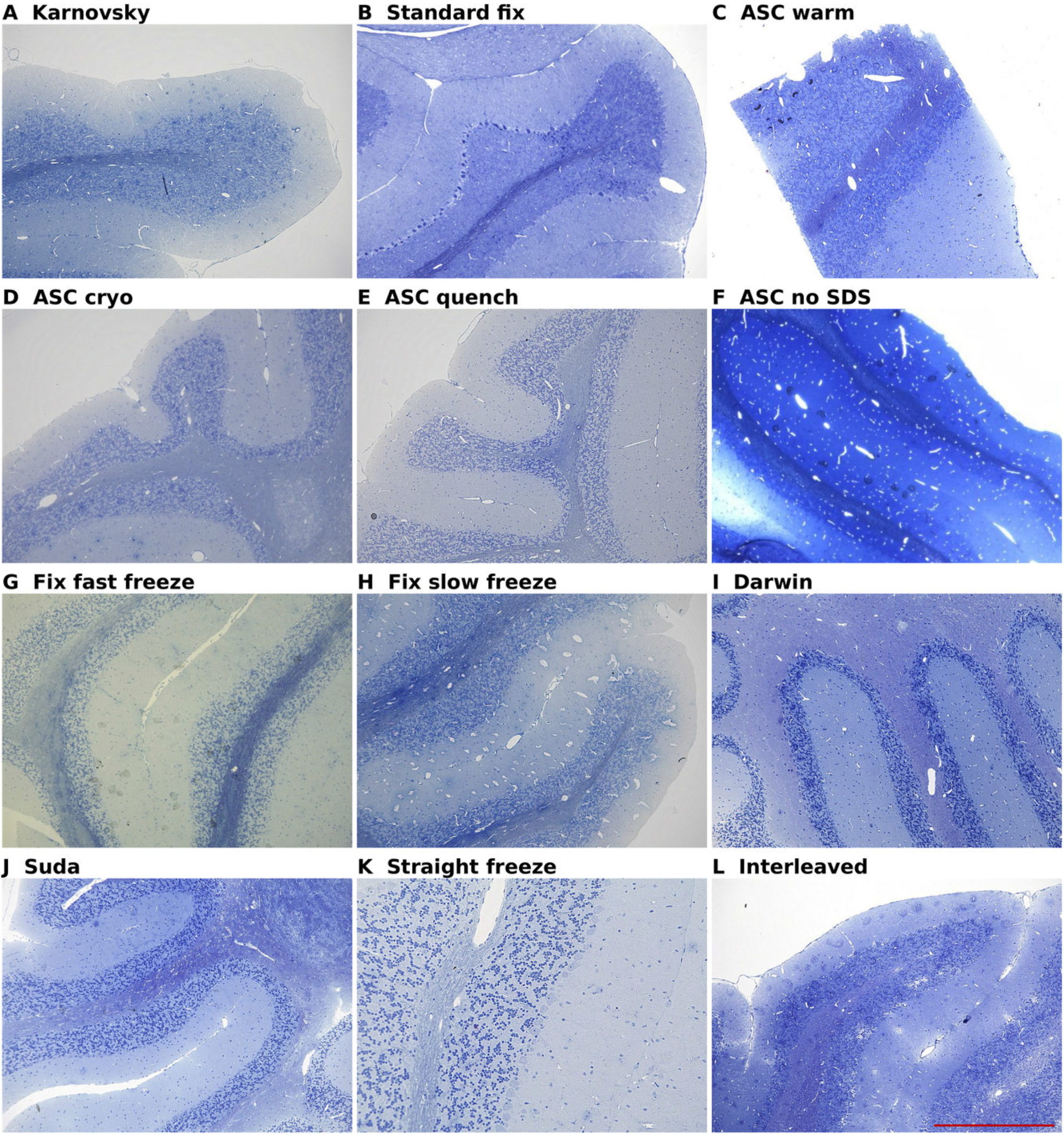
Light microscopy of cerebellar tissue across all twelve preservation conditions at 10x magnification. Toluidine blue-stained semithin sections. (A) Karnovsky fixation, (B) Standard fixation, (C) ASC warm, (D) ASC cryo, (E) ASC quench, (F) ASC without SDS, (G) Fixation followed by fast freezing, (H) Fixation followed by slow freezing, (I) High-concentration glycerol (Darwin), (J) Low-concentration glycerol (Suda), (K) Straight freeze, (L) Interleaved equilibration. Scale bar: 500 μm.

**Supplementary Figure S4.**
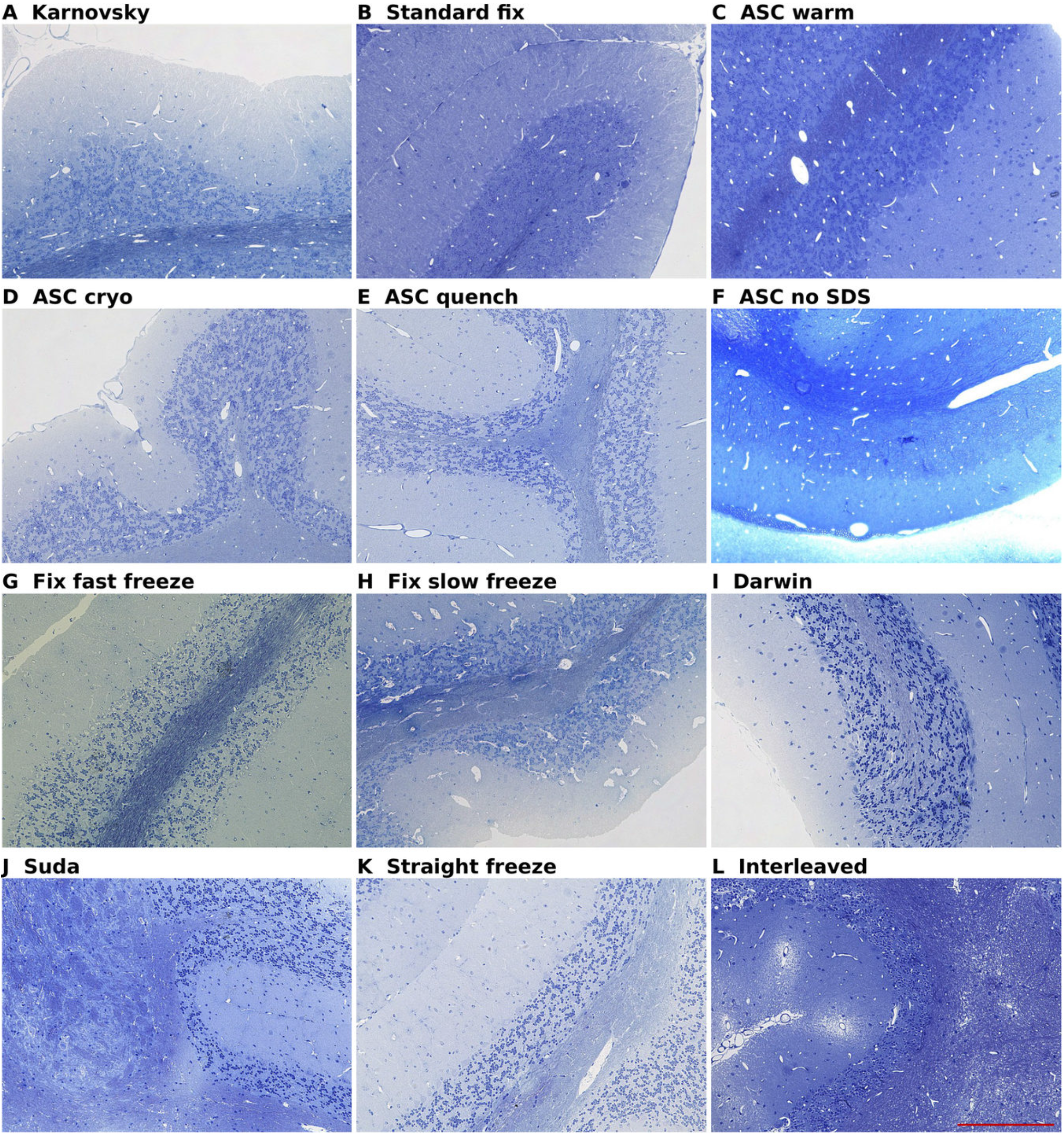
Light microscopy of cerebellar tissue across all twelve preservation conditions at 20x magnification. Toluidine blue-stained semithin sections. (A) Karnovsky fixation, (B) Standard fixation, (C) ASC warm, (D) ASC cryo, (E) ASC quench, (F) ASC without SDS, (G) Fixation followed by fast freezing, (H) Fixation followed by slow freezing, (I) High-concentration glycerol (Darwin), (J) Low-concentration glycerol (Suda), (K) Straight freeze, (L) Interleaved equilibration. Scale bar: 200 μm.

**Supplementary Figure S5.**
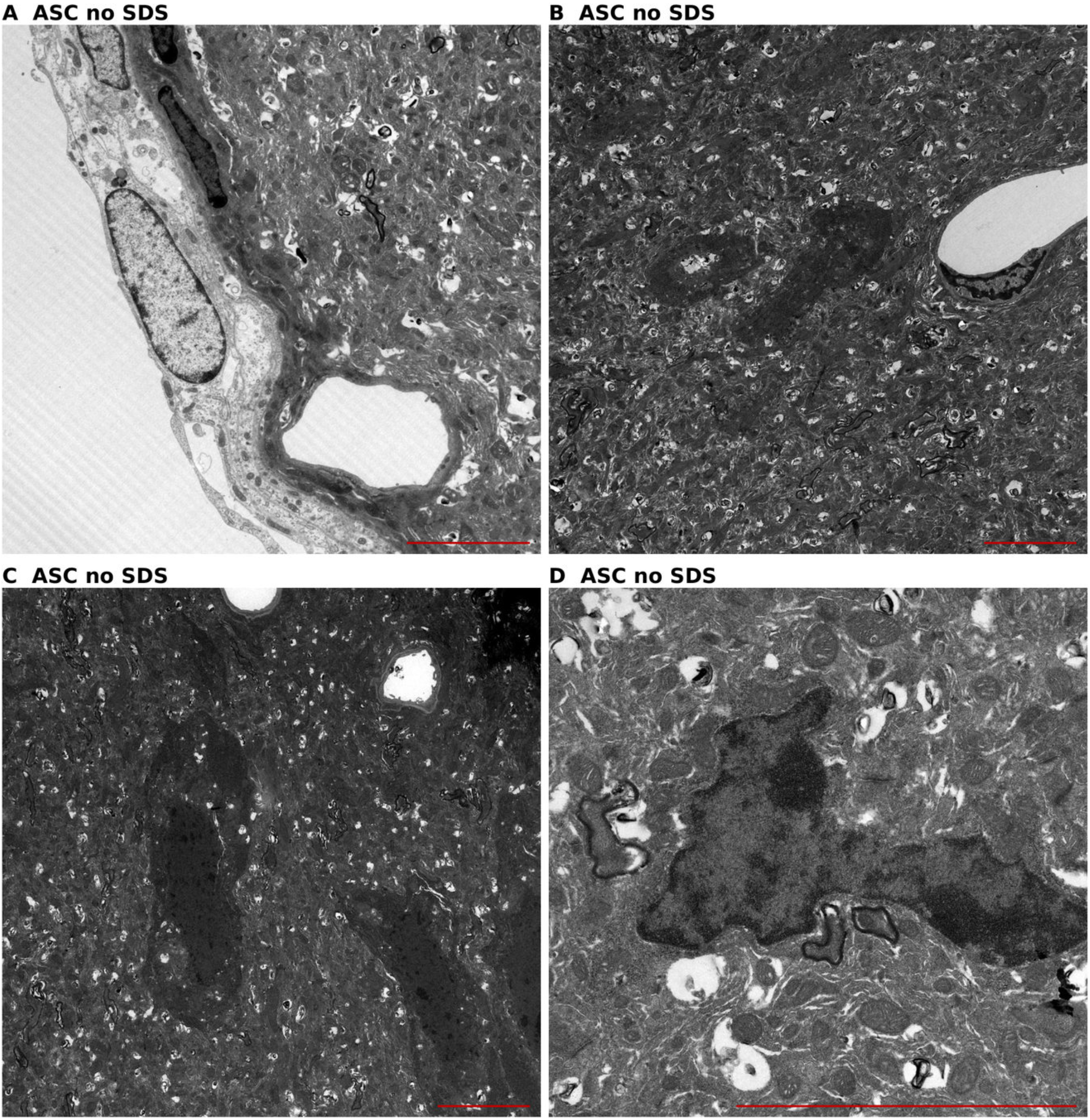
Forebrain tissue preserved with ASC without SDS. (A) Near the glia limitans, tissue appears relatively preserved with identifiable cellular structures, consistent with a superficial-to-deep gradient in water and cryoprotectant transport. (B-D) Deeper regions show increased electron density and tissue compaction consistent with dehydration from inadequate cryoprotectant penetration across the intact BBB. Scale bars: 5 μm.

**Supplementary Figure S6.**
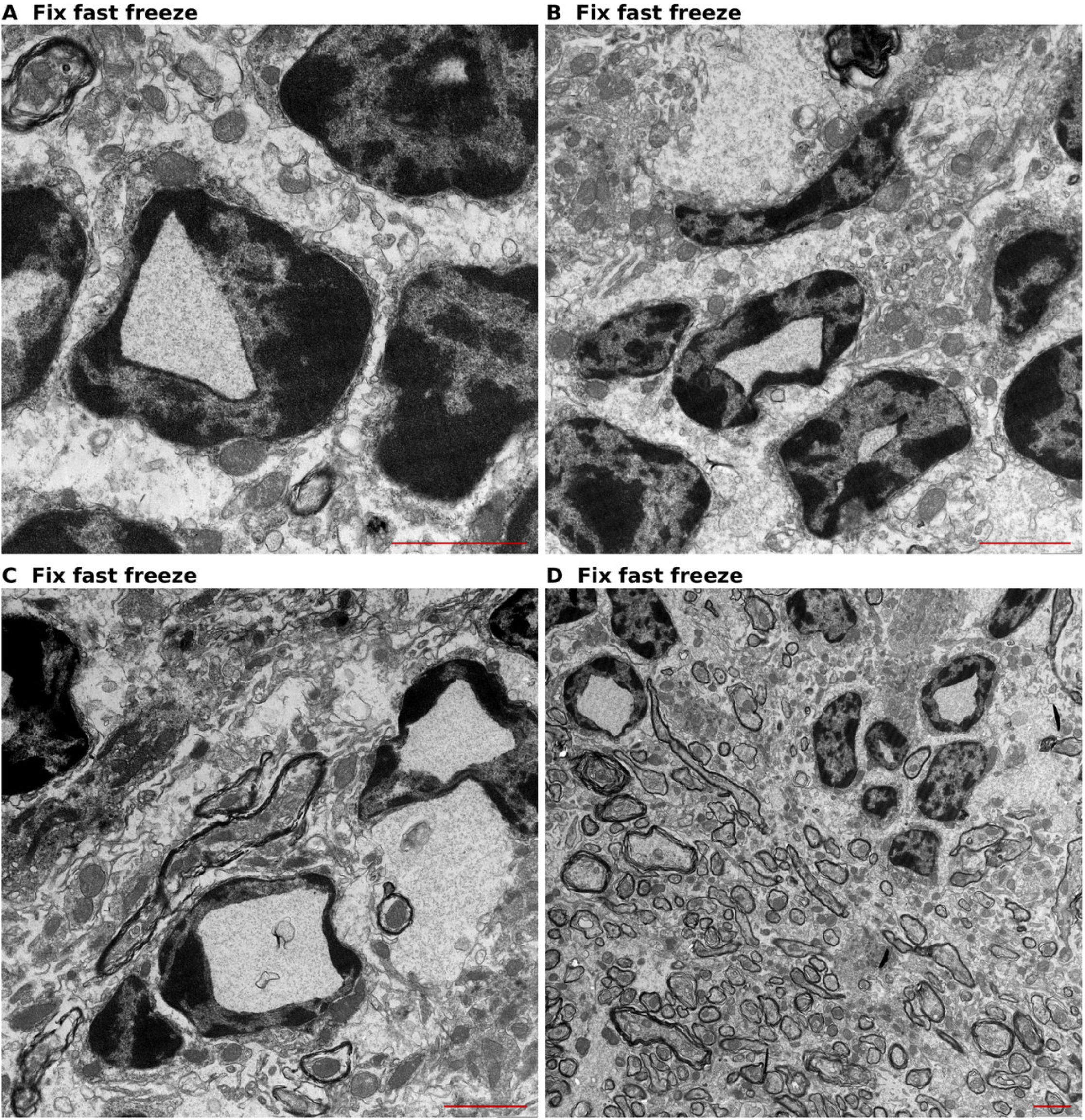
Cerebellar ultrastructure after fixation followed by fast freezing via liquid nitrogen immersion. Intranuclear cavities with rectangular and triangular geometries indicate ice crystal formation within nuclei. Scale bars: 2 μm.

**Supplementary Figure S7.**
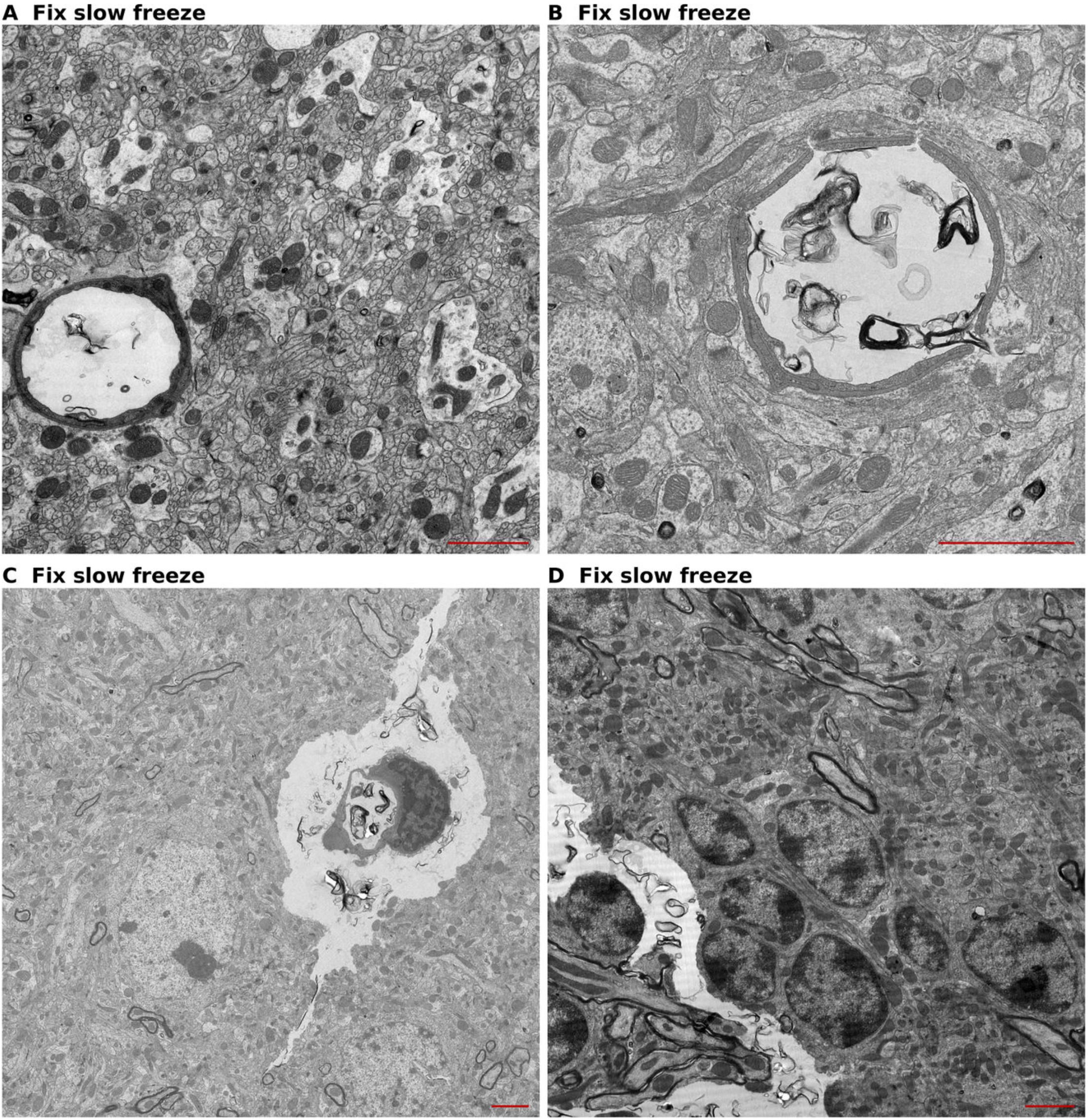
Cerebellar ultrastructure after glutaraldehyde perfusion fixation followed by slow freezing (Condition H). Perivascular microtears appear as linear clefts adjacent to capillary profiles, while parenchymal areas between vessels retain recognizable cellular structures. Scale bars: 2 μm.

**Supplementary Figure S8.**
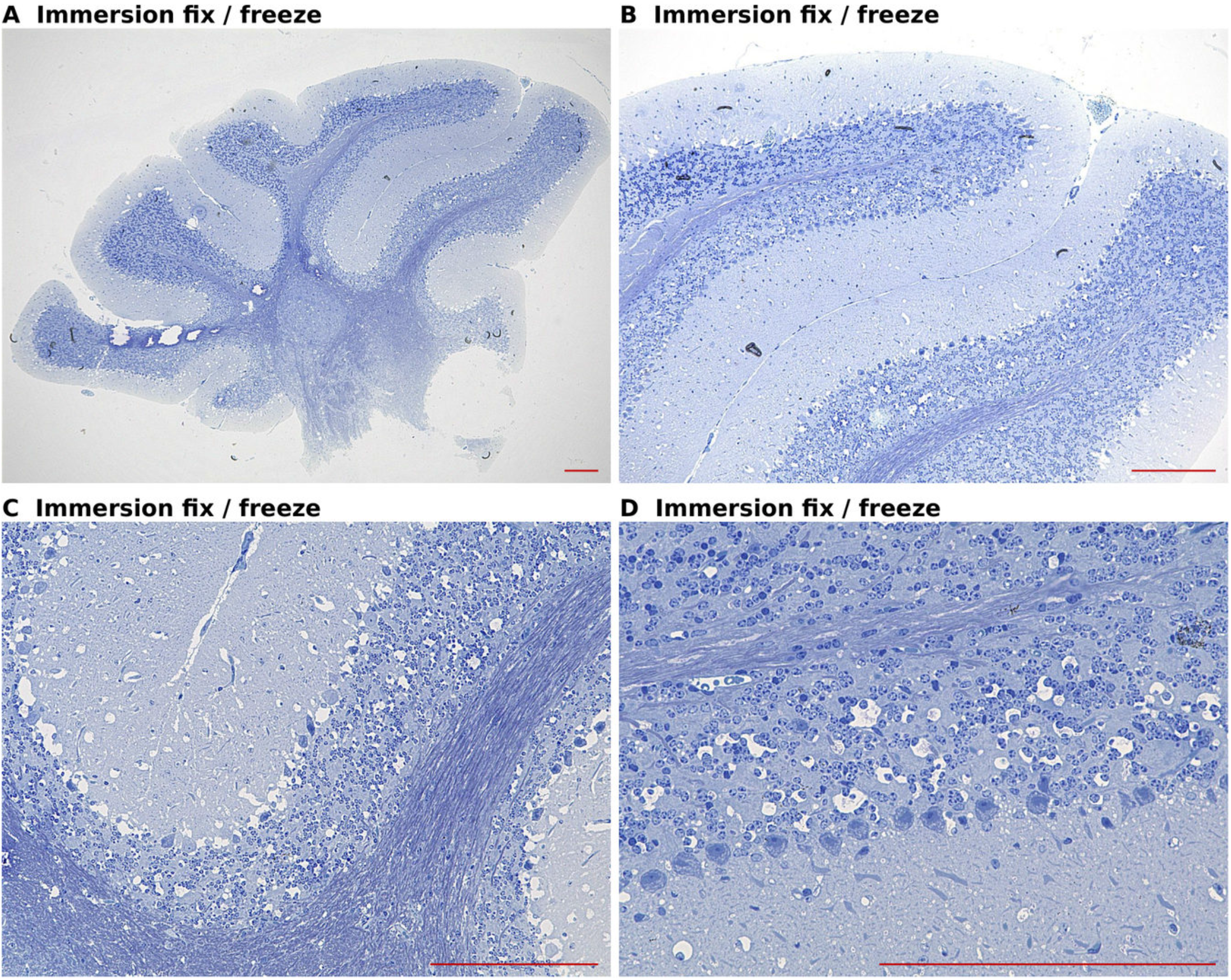
Cerebellar histoarchitecture after immersion fixation with glutaraldehyde followed by slow freezing. Microtears persist despite retention of intravascular blood contents, suggesting that the retention of intravascular material does not prevent ice-mediated mechanical disruption. Scale bar: 200 μm.

## References

1. Morton-Hayward, A.L., et al., Human brains preserve in diverse environments for at least 12 000 years. Proc. Biol. Sci., 2024. 291: p. 20232606.

2. McKenzie, A.T., et al., Cryopreservation of brain cell structure: a review. Free Neuropathol., 2024b. 5: p. 35.

3. McKenzie, A., Glutaraldehyde: A review of its fixative effects on nucleic acids, proteins, lipids, and carbohydrates. 2019. p. A.

4. McKenzie, A.T., et al., Fluid preservation in brain banking: a review. Free Neuropathol., 2024a. 5: p. 5–10.

5. Hussain, S., et al., Antibodies raised against aldehyde-fixed antigens improve sensitivity for postembedding electron microscopy. J. Neurosci. Methods, 2019. 317: p. 1–10.

6. Holleley, C.E. and E.E. Hahn, Reframing Formalin: A Molecular Opportunity Enabling Historical Epigenomics and Retrospective Gene Expression Studies. Mol. Ecol. Resour., 2025. 25: p. e14065.

7. Radzicka, A. and R. Wolfenden, Rates of Uncatalyzed Peptide Bond Hydrolysis in Neutral Solution and the Transition State Affinities of Proteases. J. Am. Chem. Soc., 1996. 118: p. 6105–6109.

8. Garrood, M., et al., Cryopreservation of aldehyde-fixed whole brains. PLoS One, 2026. 21(8): p. e0344932.

9. Laidler, K.J., Chemical Kinetics. 3rd ed. 1987, New York: Harper & Row. 44–48.

10. Fahy, G.M. and B. Wowk, Principles of Ice-Free Cryopreservation by Vitrification, in Methods Mol. 2021. p. 27–97.

11. McKenzie, A.T., et al., Biostasis: A Roadmap for Research in Preservation and Potential Revival of Humans. Brain Sciences, 2024. 14(9): p. 942.

12. Storey, K.B. and J.M. Storey, Molecular Physiology of Freeze Tolerance in Vertebrates. Physiol Rev, 2017. 97(2): p. 623–665.

13. Storey, K.B. and J.M. Storey, Biochemical adaption for freezing tolerance in the wood frog,Rana sylvatica. Journal of Comparative Physiology B, 1984. 155(1): p. 29–36.

14. Layne, J.J. and R. Jr, Freeze tolerance and the dynamics of ice formation in wood frogs (Rana sylvatica) from Southern Ohio. Canadian Journal of Zoology-revue Canadienne De Zoologie - CAN J ZOOL, 1987. 65: p. 2062–2065.

15. Costanzo, J.P., et al., Hibernation physiology, freezing adaptation and extreme freeze tolerance in a northern population of the wood frog. J Exp Biol, 2013. 216(Pt 18): p. 3461–73.

16. Polge, C., A.U. Smith, and A.S. Parkes, Revival of Spermatozoa after Vitrification and Dehydration at Low Temperatures. Nature, 1949. 164(4172): p. 666-666.

17. Smith, A.U., J.E. Lovelock, and A.S. Parkes, Resuscitation of Hamsters after Supercooling or Partial Crystallization at Body Temperatures Below 0° C. Nature, 1954. 173(4415): p. 1136-1137.

18. Lovelock, J.E. and A.U. Smith, Studies on golden hamsters during cooling to and rewarming from body temperatures below 0 degrees C. III. Biophysical aspects and general discussion. Proc R Soc Lond B Biol Sci, 1956. 145(920): p. 427–42.

19. Suda, I., K. Kito, and C. Adachi, Viability of long term frozen cat brain in vitro. Nature, 1966. 212(5059): p. 268-70.

20. Suda, I., K. Kito, and C. Adachi, Bioelectric discharges of isolated cat brain after revival from years of frozen storage. Brain Research, 1974. 70(3): p. 527–531.

21. Fahy, G.M., et al., Vitrification as an approach to cryopreservation. Cryobiology, 1984. 21(4): p. 407–26.

22. German, A., et al., Functional recovery of the adult murine hippocampus after cryopreservation by vitrification. Proceedings of the National Academy of Sciences, 2026. 123(10): p. e2516848123.

23. Pichugin, Y., G.M. Fahy, and R. Morin, Cryopreservation of rat hippocampal slices by vitrification. Cryobiology, 2006. 52(2): p. 228–240.

24. Pichugin, Y., www.cryonics.org/research/blood-brain-barrier-preliminary-patent-application-disclosure. 2007.

25. McIntyre, R.L. and G.M. Fahy, Aldehyde-stabilized cryopreservation. Cryobiology, 2015. 71: p. 448–458.

26. Meissner, D.H. and H. Schwarz, Improved cryoprotection and freeze-substitution of embryonic quail retina: A tem study on ultrastructural preservation. Journal of Electron Microscopy Technique, 1990. 14(4): p. 348–356.

27. Fahy, G.M., et al., Ultrastructural and Histological Cryopreservation of Mammalian Brains by Vitrification. 2026.

28. Darwin, M., et al., Effect of Human Cryopreservation Protocol on the Ultrastructure of the Canine Brain, in CryoNet. 1995.

29. Ito, S., Formaldehyde-glutaraldehyde fixatives containing tri nitro compounds. J Cell Biol, 1968. 39: p. 168A–169A.

30. German, A., Perfusion quenching. Cryobiology, 2024. 117: p. 105097.

31. Oquab, M., et al., DINOv2: Learning Robust Visual Features without Supervision. 2024. p. Learning.

32. Dong, W., C. Moses, and K. Li. Efficient k-nearest neighbor graph construction for generic similarity measures. in Association for Computing Machinery, New York, NY, USA, pp. 2011. Association for Computing Machinery.

33. Rubinsky, B. and D.E. Pegg, A mathematical model for the freezing process in biological tissue. Proc. R. Soc. Lond. B Biol. Sci., 1988. 234: p. 343–358.

34. Schmitt, A., et al., Optimized protocol for whole organ decellularization. Eur J Med Res, 2017. 22: p. 31.

35. Butt, A.M., H.C. Jones, and N.J. Abbott, Electrical resistance across the blood-brain barrier in anaesthetized rats: a developmental study. J Physiol, 1990. 429: p. 47–62.

36. MacAulay, N., Molecular mechanisms of brain water transport. Nat Rev Neurosci, 2021. 22(6): p. 326–344.

37. Manley, G.T., et al., Aquaporin-4 deletion in mice reduces brain edema after acute water intoxication and ischemic stroke. Nat Med, 2000. 6(2): p. 159–63.

38. Herscovitch, P., et al., Positron Emission Tomographic Measurement of Cerebral Blood Flow and Permeability—Surface Area Product of Water Using [15O]Water and [11C]Butanol. Journal of Cerebral Blood Flow & Metabolism, 1987. 7(5): p. 527–542.

39. Ahmadkhani, N., et al., High throughput method for simultaneous screening of membrane permeability and toxicity for discovery of new cryoprotective agents. Scientific Reports, 2025. 15(1): p. 1862.

40. Tewes, B.J. and H.J. Galla, Lipid polarity in brain capillary endothelial cells. Endothelium, 2001. 8: p. 207–220.

41. Gurtovenko, A.A. and J. Anwar, Modulating the structure and properties of cell membranes: the molecular mechanism of action of dimethyl sulfoxide. J. Phys. Chem. B, 2007. 111: p. 10453–10460.

42. Trump, B.F., et al., Effects of Freezing and Thawing on the Structure, Chemical Constitution, and Function of Cytoplasmic Structures. Fed. Proc., 1965. 24: p. S144–168.

43. Ngapo, T.M., et al., Freezing rate and frozen storage effects on the ultrastructure of samples of pork. Meat Sci., 1999. 53: p. 159–168.

44. Menz, L.J., Structural changes and impairment of function associated with freezing and thawing in muscle, nerve, and leucocytes. Cryobiology, 1971. 8(1): p. 1–13.

45. Oehmichen, M. and M. Gencic, Postmortal diffusion of plasma albumin in rat brain. Zeitschrift für Rechtsmedizin, 1980. 84(2): p. 113–123.

46. Moen, O.M., The case for cryonics. J Med Ethics, 2015. 41(8): p. 677–81.

